# Neuroligin-2 is ubiquitinated by Nedd4l to control developmental astrocyte morphogenesis

**DOI:** 10.64898/2025.12.15.694023

**Authors:** Kristina Sakers, Juan J. Ramirez, Nimrod Elazar, Leykashree Nagendren, Erik Soderblom, Cagla Eroglu

## Abstract

Central nervous system astrocytes have an intricate, highly branched morphology. Proper development of perisynaptic astrocyte processes is necessary for tripartite synapse formation and function. However, cellular pathways orchestrating this development are largely unknown. Neuroligins (NLs) 1-3 regulate astrocyte morphogenesis via transcellular adhesions with neuronal neurexins. Here, we found an astrocytic NL2-based mechanism governing morphogenesis. Through structure and function studies, we identified a WW-binding motif within the NL2 intracellular domain required for astrocyte morphogenesis. Using cell-specific *in vivo* proximity labeling (iBioID), we found that each NL displays distinct protein-protein interactions within astrocytes, distinct from the neuronal NL2 binding partners. From these data, we identified a role for WW domain-containing E3 ubiquitin ligase Nedd4l in astrocyte morphogenesis. Biochemical assays revealed Nedd4l ubiquitinates and stabilizes NL2, and this ubiquitination is required for astrocyte morphogenesis. This study shows that Neuroligins have non-overlapping roles in controlling astrocyte growth and uncovers a molecular mechanism of how NL2 mediates astrocyte morphogenesis.

**SUMMARY:** Sakers et al report that astrocytic neuroligins (NLs) are functionally diverse proteins, with unique intracellular protein-protein interactions. They show that Nedd4l binds to NL2 and ubiquitinates its intracellular domain, leading to changes in NL2 stability. In vivo, Nedd4l and NL2 ubiquitination are critical for astrocyte morphogenesis.

## INTRODUCTION

Protoplasmic astrocytes of the central nervous system (CNS) have a unique bush-like morphology. The molecular mechanisms that govern how astrocytes develop their complex structure are largely unknown. Astrocytes tile the brain and extend fine filopodia-like processes into the neuropil (Bushong et al., 2004, 2002), facilitating an active role in critical brain functions, including synaptogenesis, regulation of synaptic strength, and buffering of neurotransmitters and ions (Farizatto and Baldwin, 2023; Verkhratsky and Nedergaard, 2018). Human astrocytes are disproportionately larger and more complex than rodent astrocytes (Oberheim et al., 2006), suggesting that higher-order species require astrocytes for advanced brain function. Consequently, astrocyte dysfunction has been identified across a myriad of neurological diseases (Tian et al., 2010; Rothstein et al., 1995; Preman et al., 2021; Booth et al., 2017; Gutmann et al., 1999). In particular, a recent study found that numerous Alzheimer’s risk genes correlate specifically with astrocyte territory size (Endo et al., 2022). Thus, it is crucial to understand how astrocyte morphogenesis proceeds and which cell biological mechanisms govern this process.

Various signals have emerged as regulators of astrocyte growth, including the neuron-derived secreted factors Sonic Hedgehog (Xie et al., 2022) and BDNF (Holt et al., 2019), actin-binding proteins such as Ezrin (Endo et al., 2022; Lavialle et al., 2011; Wang et al., 2025) and Profilin-1 (Molotkov et al., 2013), and numerous cell-adhesion molecules (Baldwin et al., 2023). Among the latter, Neuroligins (NLs) play a critical role in the morphogenesis of astrocytes (Ackerman et al., 2021; Stogsdill et al., 2017), as well as oligodendrocyte precursor cells (Li et al., 2024). NLs are trans-synaptic cell-adhesion proteins that bind to Neurexins to promote synaptic formation (Barrow et al., 2009; Chih et al., 2005; Scheiffele et al., 2000; Shipman and Nicoll, 2012) and function (Babaev et al., 2016; Varoqueaux et al., 2006). There are four NL genes expressed in rodents, with NL1, NL2, and NL3 being the predominantly expressed family members. In rodent neurons, NL1 dose-dependently controls dendritic morphology, with NL1 overexpression increasing and NL1 knockdown decreasing dendritic arborization (Schnell et al., 2014, 2012). In contrast, loss of NL3 results in a more complex dendritic arbor (Xu et al., 2019), suggesting non-overlapping roles for NLs in controlling neuronal morphology. Neuronal NLs also have non-redundant roles in controlling synaptic function (Chanda et al., 2017; Nguyen et al., 2016). In astrocytes, loss of NLs 1-3 each reduces astrocyte territory and neuropil infiltration size (Stogsdill et al., 2017). However, whether NLs play unique roles in astrocyte morphogenesis and what mechanisms control this process is unclear. NLs are implicated in neurodevelopmental diseases, including Autism Spectrum Disorder (Jamain et al., 2003; Parente et al., 2017), and thus, it is imperative to understand NL function across all cell types throughout development.

Functional differences across NLs may stem from amino acid sequence divergence. Rodent NLs 1-3 have similar extracellular domains (ECD, ∼75% homology) and more disparate intracellular domains (ICD, ∼50% homology). Several amino acid motifs in NL ICDs have been shown to direct interactions with specific protein partners, ultimately leading to unique functional consequences such as the specification of inhibitory versus excitatory synapses (Poulopoulos et al., 2009; Shipman et al., 2011; Sumita et al., 2007). One study using chimeric NLs demonstrated that NL function is not solely dependent on Neurexin binding to the ECD, but also requires interactions with intracellular proteins to control synapse-specific functions (Nguyen et al., 2016). Further, post-translational modifications on NL ICDs can influence protein-protein interactions and downstream signaling. For example, NL1 tyrosine phosphorylation at Y782 results in preferential binding to the excitatory postsynaptic protein PSD-95 over the inhibitory postsynaptic scaffold protein Gephyrin. PSD95 then recruits α-amino-3-hydroxy-5-methyl-4-isoxazolepropionic acid (AMPA) receptors and controls synaptic firing (Giannone et al., 2013; Letellier et al., 2018). These studies have been conducted in a neuronal context; thus, whether astrocytic NLs exhibit distinct protein-protein interactions and how these interactions control astrocyte development remain unknown.

In this study, we show that NLs 1-3 are not redundant in controlling astrocyte morphological development. We identify that the WW-binding motif in NL2 is necessary for astrocyte morphogenesis. Using an unbiased *in vivo* proximity-labeling approach, we uncover interaction networks for astrocytic NL1, NL2, and NL3. We also performed proximity labeling of neuronal NL2 to compare cell-type-specific differences in its interactome and function. We show that NLs have distinct interactomes representing unique functions within astrocytes and between astrocytes and neurons. These data reveal numerous WW-domain-containing proteins that are putative interactors of WW motifs in NLs. Of these, we identified Nedd4l, a HECT-domain E3 ubiquitin ligase, to control NL2 stability via ubiquitination, which is necessary for astrocyte morphogenesis. Moreover, the loss of Nedd4l in astrocytes results in a stunted astrocyte morphology. Importantly, Nedd4l interacts with NL1 and NL2, but not NL3. Together, this study reveals unique interactions of astrocytic NL2, providing a previously unknown mechanism for astrocyte morphological development.

## RESULTS

### Astrocytic Neuroligins 1, 2, and 3 have non-redundant roles in morphogenesis

NLs 1, 2, and 3 are all expressed throughout development in mouse cortical astrocytes (Boisvert et al., 2018; Clarke et al., 2018; Zhang et al., 2014; Farhy-Tselnicker et al., 2021). Loss of either NL1, NL2, or NL3 from cortical astrocytes significantly reduces astrocyte complexity, and the loss of all three NLs together results in an additive reduction of complexity (Stogsdill et al., 2017), indicating that these proteins play distinct roles during astrocyte development. NLs have similar ECD amino acid sequences yet differ substantially in their intracellular domains (Fig 1A). Given differences in amino acid sequences and the combinatorial deficit in astrocyte complexity, we examined whether NLs have redundant roles in astrocyte morphogenesis. To do so, we used an optimized and validated *in vitro* astrocyte-neuron co-culture method to quantify astrocyte branch complexity via Sholl analysis (Stogsdill et al., 2017; Baldwin et al., 2021; Tan et al., 2023) (Fig 1B). In this *in vitro* system, astrocytes require neuronal contact to develop a highly branched morphology (Baldwin et al., 2021; Stogsdill et al., 2017) and we can manipulate astrocyte gene expression without affecting the neurons. Because NL2 loss affects astrocyte morphogenesis *in vivo* at both early and late developmental time points, we focused on this NL. We transfected astrocytes with a plasmid expressing an shRNA for *Nlgn2* (shNL2), which has been validated for NL2 knockdown efficiency and NL2 specificity (Fig S1A, B) and extensively tested for off-target effects (Stogsdill et al., 2017; Chih et al., 2006; Singh et al., 2016). This plasmid also encodes CAG-GFP to visualize transfected cells. Loss of *Nlgn2* in astrocytes causes a substantial reduction in astrocyte morphology when co-cultured with neurons (Fig S1C-D; Fig 1C-D). To provide additional evidence for the role of Nlgn2 in astrocyte morphogenesis and to address potentially confounding effects of shRNA toxicity (Goel and Ploski, 2022), we designed and tested gRNAs targeting rat *Nlgn2* (Fig S1E). Two of three of the gRNAs targeting *Nlgn2* caused a translation-disrupting frameshift mutation, reflected by the loss of GFP expression downstream of the gRNA and PAM sequences (see Methods) (Fig S1F). We expressed these gRNAs in primary rat astrocytes, along with *Staphylococcus aureus* Cas9 (SaCas9) (Hunker and Zweifel, 2020), under the control of the GfaABC1D promoter (Lee et al., 2008) and found that both gRNAs significantly reduced the morphological complexity in transfected astrocytes cultured with WT neurons, compared to the empty vector control (Fig S1G, H). These data further support that the morphological phenotype observed in Nlgn2 shRNA-transfected cells cannot be explained by cytotoxicity.

**Figure 1:**
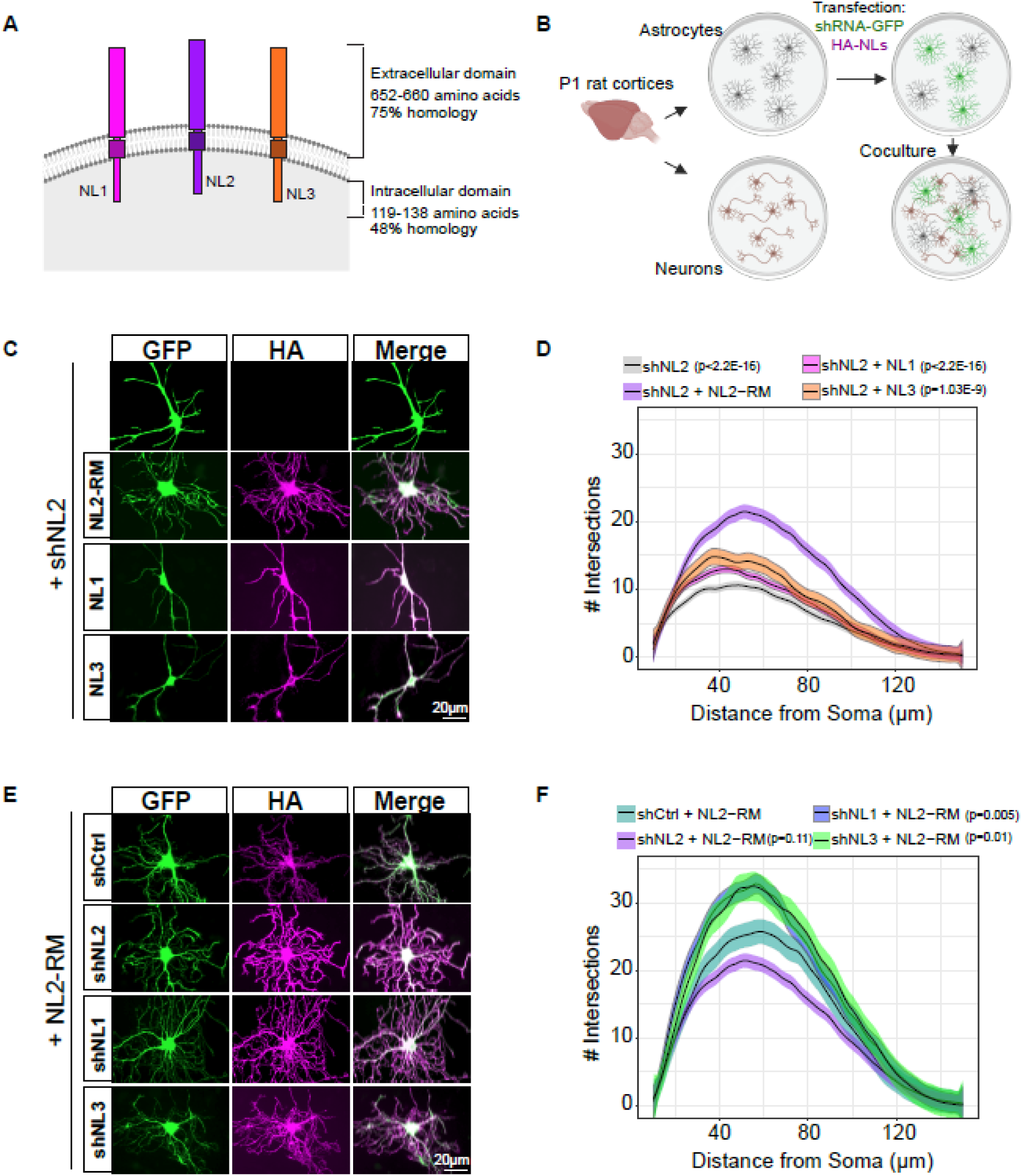
Astrocytic Neuroligins 1, 2, and 3 have non-redundant roles in morphogenesis. A) Cartoon depiction of NLs 1-3. Length and homology refer to mouse sequences. B) Neuron-astrocyte co-culture method to study astrocyte morphogenesis in vitro. P1 rat cortices are used to isolate astrocytes and neurons separately to allow for astrocyte-specific gene manipulation. Astrocytes post-transfection are plated onto neurons for 48h to allow arborization. C) Representative images of astrocytes transfected with shNL2 and either: HA-NL2-RM (Rescue mutant, resistant to shRNA), HA-NL1, or HA-NL3. Scale bars are 20µm. D) Sholl analysis quantification of conditions shown in C. Linear-mixed model followed by ANOVA reveals main effect of condition F(3, 319) = 42.46, p < 2.2E-16. P-values represent Dunnett’s post-hoc with respect to the shNL2+HA-NL2-RM condition. N = 48-96 cells per condition, across >3 independent culture experiments. See Figure S1C-D for a comparison of shNL2 to shCtrl astrocytes. Shaded area indicates standard error. E) Representative images of astrocytes co-transfected with shRNA against either NL1, NL2, or NL3, and HA-NL2-RM. Scale bars are 20µm. F) Sholl analysis quantification of conditions in E. Linear-mixed model followed by ANOVA reveals trending effect of condition F(3, 311)=15.5, p=2.02E-9. P-values represent Dunnett’s post-hoc with respect to shCtrl+NL2-RM. N = 53-97 cells per condition, across 3 independent culture experiments. Note that the shNL2 + HA-NL2 data is the same across D and F. Shaded area indicates standard error.

Next, we tested whether the stunted morphological phenotype of *Nlgn2* knockdown astrocytes can be rescued by the co-expression of an HA-tagged NL2 that has been modified to be shNL2 resistant (HA-NL2-RM, Fig S1A, B) (Stogsdill et al., 2017) or via the co-expression of HA-tagged NL1 or NL3 with shNL2, to investigate whether increasing total NL levels can compensate for the loss of another. Interestingly, HA-tagged NL1 or NL3 did not rescue the astrocyte morphology (Fig 1C, D). In contrast, co-expression of HA-NL2 with either shNL1 or shNL3 resulted in a higher number of maximum intersections in the astrocyte branches compared to a scrambled shRNA control (shCtrl) (Fig 1E, F). Together, these data show that NLs 1, 2, and 3 do not have redundant roles in controlling astrocyte morphogenesis. Moreover, these findings suggest that NL2 is indispensable in astrocyte morphological development.

### The intracellular WW-binding motifs of Neuroligins uniquely regulate astrocyte morphology

Previous work has revealed unique roles for NL ICDs in controlling synaptic transmission (Nguyen et al., 2016). Thus, we hypothesized that the ICDs of NLs may underlie their distinct roles in controlling astrocyte morphogenesis. NLs contain multiple protein motifs in their ICDs (Fig S2A) that regulate binding to synaptic scaffolding molecules, including Gephyrin and Collybistin (Poulopoulos et al., 2009), which are necessary for inhibitory synapse specification, and PSD-95 (Irie et al., 1997), which organizes excitatory postsynaptic sites. We wondered if any of these motifs in the NL2 ICD are necessary for astrocyte morphology. We used mutagenesis to either: delete the NL2 PDZ binding domain (NL2 ΔPDZ, deletion of amino acids 833-836), delete the NL2 polyproline-rich motif, which activates Collybistin (NL2 ΔPRM, deletion of amino acids 798-805) or mutate NL2 Gephyrin binding residue (NL2 GBM, Y770A) (Fig 2A). All of these constructs use the HA-NL2-RM sequence to prevent their degradation by shNL2. We tested whether these mutant constructs could rescue the shNL2 morphological deficits in our co-culture system. These mutants were properly trafficked to the cell surface (Fig S2B) and rescued the morphological deficits in shNL2 astrocytes (Fig 2B-C), suggesting that potential interactions with Gephyrin, Collybistin, and PSD-95 are dispensable for NL2 control of astrocyte morphology.

**Figure 2:**
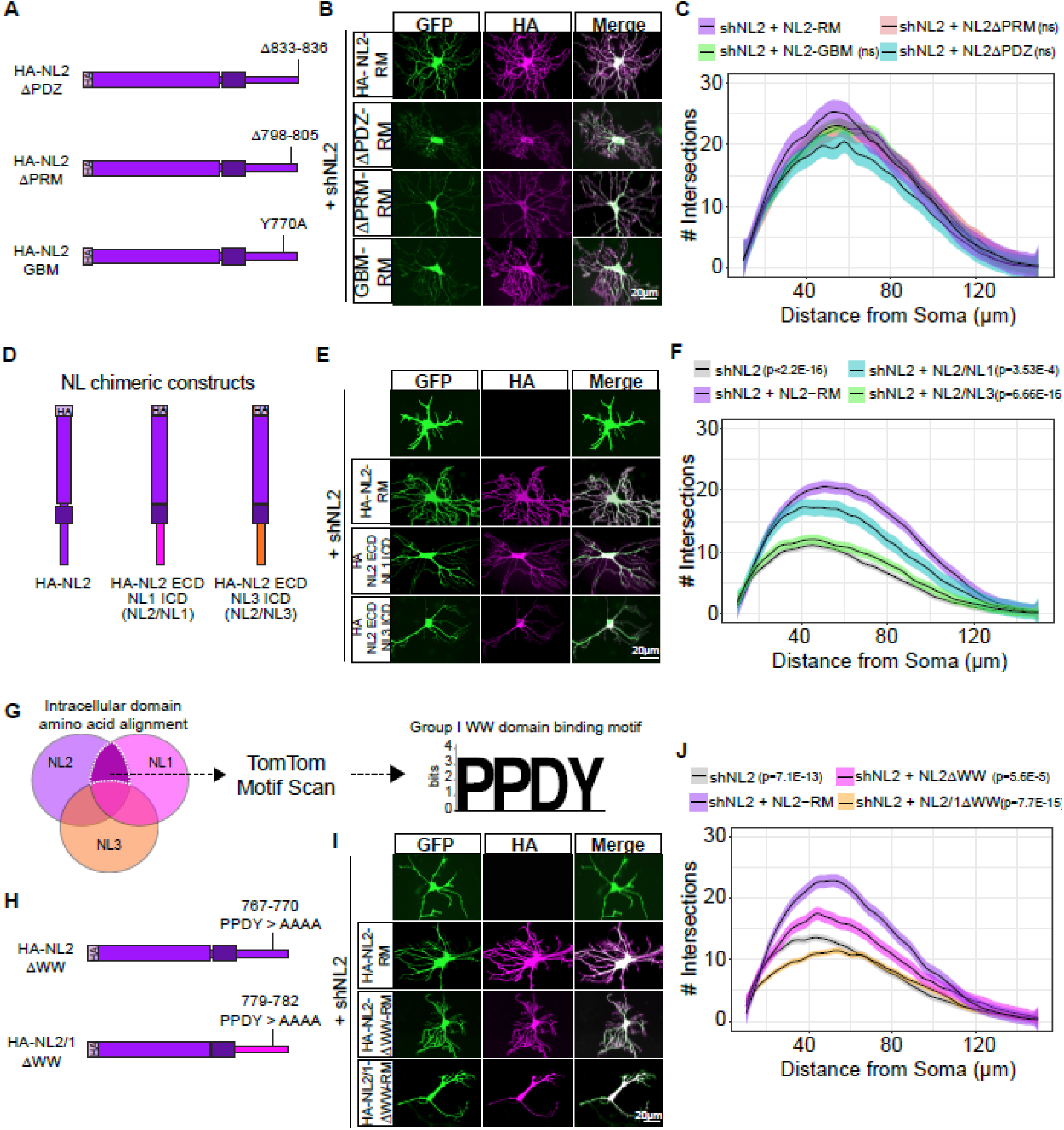
The intracellular WW-binding motifs of Neuroligins uniquely regulate astrocyte morphology. A) Cartoon of NL2 intracellular domain mutants. B) Representative images of astrocytes co-transfected with shNL2 and either HA-NL2-RM (Rescue Mutant, resistant to shRNA) or mutants lacking known motifs. Scale bars are 20µm. C) Sholl analysis quantification of the conditions in D. Linear-mixed model followed by ANOVA reveals no effect of condition F(3, 159)=1.97, p=0.12. NS = not significant in Dunnett’s post-hoc. N=36-46 cells per condition, across 3 independent culture experiments. Shaded area indicates standard error. D) Cartoon of NL2-ECD and NL1 or 3 ICD chimeric constructs. E) Representative images of astrocytes transfected with shNL2 and either HA-NL2-RM (Rescue mutant, resistant to shRNA), or chimeric proteins: HA-NL2(ECD)-NL1(ICD), HA-NL2(ECD)-NL3(ICD). Scale bars are 20µm. F) Sholl analysis quantification of conditions in B. Linear-mixed model followed by ANOVA reveals the main effect of condition F(3,356)=38.22, p<2.2E-16. P-values represent Dunnett’s post-hoc with respect to shNL2+NL2-RM. N=57-97 cells per condition, across 4 independent culture experiments. Shaded area indicates standard error. G) Venn diagram depicting the overlap of sequences in NL1, NL2, and NL3 ICDs used for motif scanning by TomTom. We identified the motif PPDY which is common between NL1 and NL2 but absent in NL3. H) Cartoon of NL2ΔWW and NL2/1ΔWW mutant constructs. I) Representative images of astrocytes transfected with shNL2 and HA-NL2-RM-ΔWW or shNL2 and HA-NL2/1-ΔWW. Scale bars are 20µm. J) Sholl analysis quantification of conditions in F. Linear-mixed model ANOVA reveals the main effect of condition F(3, 304)=25.76, p=7.1E-15. P-values represent Dunnett’s posthoc with respect to shNL2+NL2-RM. N = 71-84 cells per condition, across 4 independent culture experiments. Shaded area indicates standard error.

To identify the regions of the NL2 ICD that are required for astrocyte morphogenesis, we used previously validated (Nguyen et al., 2016) NL2 chimeric constructs to investigate whether replacing the NL2 ICD with either the NL1 or the NL3 ICD is sufficient to rescue the loss of NL2 in astrocytes (Fig 2D). Using astrocyte-neuron co-cultures, we expressed the NL chimeras (HA-NL2-RM sequence) in astrocytes in the presence of shNL2 and quantified astrocyte complexity (Fig 2E). While the NL2 ECD/NL1 ICD condition resulted in more complex astrocytes than shNL2 alone, the astrocytes transfected with NL2 ECD/NL3 ICD were identical to shNL2 alone (Fig 2F). This finding suggests that the NL1 and NL2 ICDs share a common amino acid sequence required for controlling astrocyte morphology, which is absent from the NL3 ICD.

To pinpoint the interaction motif in the NL2 ICD that is required for astrocyte morphogenesis, we aligned the intracellular domain amino acid sequences of NL1-3 (Fig 2G, Fig S2A) and isolated motifs that were present in both NL1 and NL2, but absent from NL3. Using TomTom motif scanning (Gupta et al., 2007), we found the sequence PPDY – a Group I WW domain binding motif, known to have the consensus sequence of PPxY (Kasanov et al., 2001; Pirozzi et al., 1997), to fit our criterion. When we mutated the PPDY motif of HA-NL2-RM to four alanines, we found that this construct could not rescue the astrocyte complexity (Fig 2H-J). Similarly, mutation of the PPDY (PPDY > AAAA) in the NL2 ECD/NL1 ICD chimera ablated the morphogenesis rescue previously seen for the non-mutated chimera (Fig 2H-J). Together, these data suggest that NL2 controls astrocyte morphogenesis through PPDY motif interactions in its ICD.

### *In vivo* BioID reveals unique interactions of astrocytic Neuroligins, including WW domain-containing proteins

PPxY motifs are recognized by Group I WW domain-containing proteins (Kato et al., 2004, 2002). WW domains are ∼40 amino acids, with two flanking conserved tryptophan (single amino acid code: W) residues (Sudol and Hunter, 2000). In the human proteome, there are over 50 proteins that contain a WW domain, with many containing multiple WW domains. To identify which WW domain-containing protein binds NL1 and NL2, we used *in vivo* BioID (iBioID). This approach (Roux et al., 2012) uses a biotin ligase, Turbo BirA, fused to a bait protein to identify all proteins interacting with the bait, within 10nm (Branon et al., 2018). We made C-terminal fusions of Turbo BirA to NL1, 2, or 3 and virally expressed these constructs under the control of the astrocyte promoter GfaABC1D (Lee et al., 2008) (Fig 3A-D). To investigate cell-type specific differences in the NL2 interactome, we also expressed NL2-Turbo BirA under the control of the neuronal promoter hSyn1 (Fig 3A-B). As controls, we expressed soluble TurboID in astrocytes or neurons, which will biotinylate the repertoire of intracellular proteins in each cell-type. We chose to use a cytosol-localized control, as NLs are trafficked throughout many cellular compartments (Kang et al., 2014; Binda et al., 2019; Halff et al., 2019; Xie et al., 2023; Xu et al., 2024), and we aimed to capture all physiologically relevant interactors, not just those at the plasma membrane. We confirmed that the NL2-Turbo BirA constructs were properly localized to the cell membrane in cultured astrocytes (Fig S3A) and they could rescue the morphological deficits of shNL2 astrocytes (Fig S3B, C), thus indicating that NL2 functioned as expected. However, we found that placing the Turbo BirA after the PDZ-binding motif in the NL2 ICD reduced the enzyme’s biotinylation activity. By moving the PDZ domain after the TurboID, we observed robust biotinylation (Fig S3B) and proceeded with this placement of the TurboID (before the PDZ binding motif) for all NL constructs (Fig 3A).

**Figure 3:**
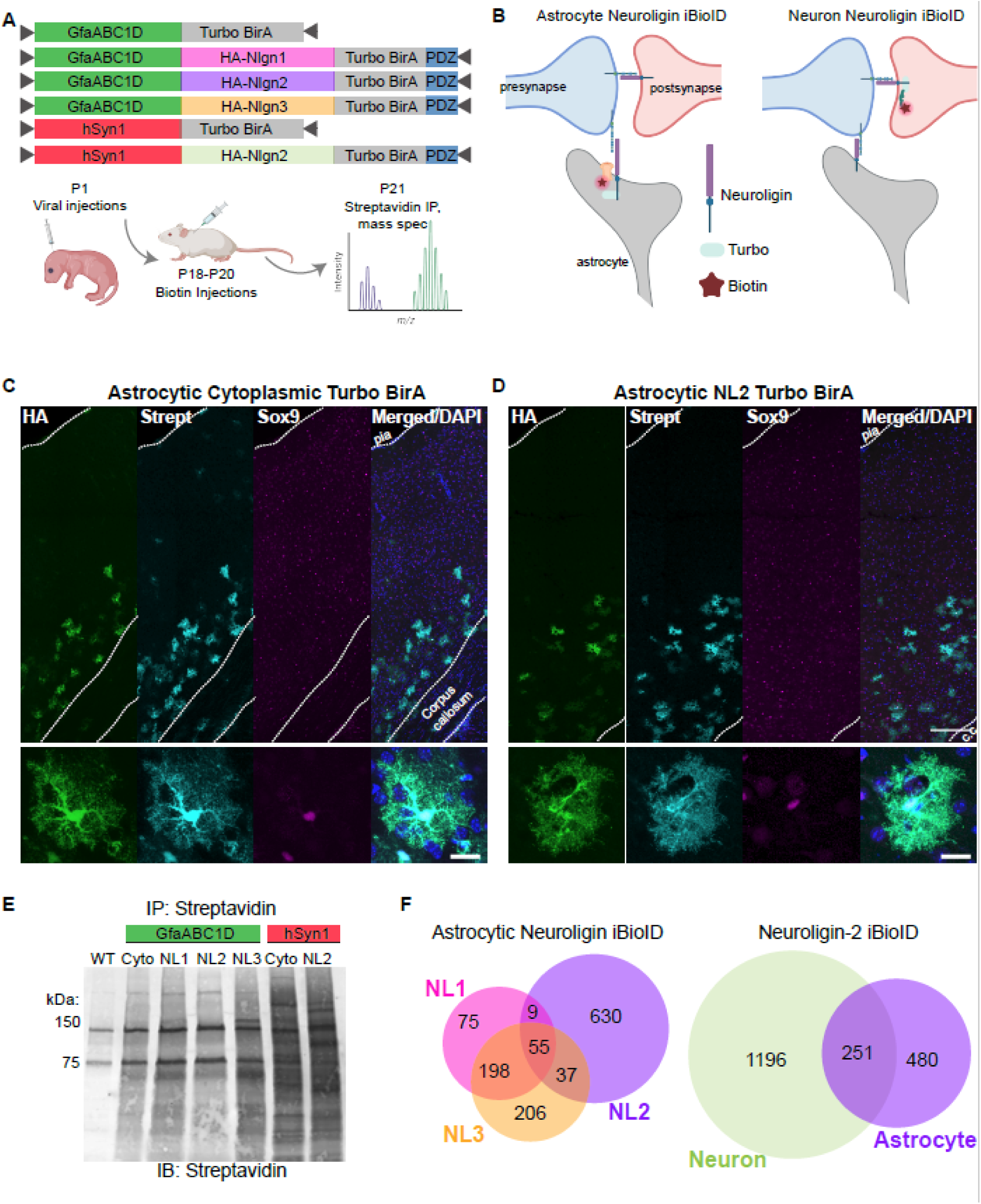
*In vivo* Neuroligin BioID. A) Schematic of AAV constructs and the experimental strategy of in vivo BioID to identify NL interactome *in vivo*. Each biological replicate contained two pooled cortices, 3 biological replicates per condition. B) Cartoon depiction of Neuroligin iBioID constructs in astrocytes or neurons. C-D). Representative cortical images of Astrocytic Turbo BirA (C) or Astrocytic NL2 Turbo BirA (D) injected animals. Stained with Streptavidin-594 (Strept), HA and Sox9 to determine astrocyte specificity. Scale bars: column image = 100µm; inset = 20µm. E) Representative western blot of iBioID immunoprecipitation from astrocytic Turbo vs NL2-Turbo animals demonstrates successful enrichment of biotinylated proteins in immunoprecipitation. Input is 2% of total cortex homogenate. IP is 15% of the eluate sample. Prominent bands are expected at approximately 150 and 75kDa which represent mitochondrial carboxylases(Kirkeby et al., 1993) [blot is overexposed to show this in the input lanes]. Resulting ‘smear’ in IP lanes indicated various-sized proteins that have been biotinylated near the bait protein. F) Venn diagram of protein interactors across astrocytic NL1, NL2, and NL3 and astrocytic versus neuronal NL2.

We injected perinatal mice (P0-P1) intracortically with AAVs expressing the NL-Turbo BirA fusions, or the cell-type-specific soluble BirA control. After three days of biotin injection (P18-P20), we collected P21 mouse cortices and isolated biotinylated proteins by streptavidin immunoprecipitation (Figs 3A, E). We chose P21 as an endpoint because NL2 loss in developing astrocytes results in significantly smaller astrocyte territories with concomitant deficits in synapse formation and function at P21 (Stogsdill et al., 2017). We confirmed the cell-type-specificity of biotinylation *in vivo* by co-staining for the astrocytic marker Sox9 (Fig 3C-D) or the neuronal marker NeuN (Fig S3D). Mass spectrometry identification and quantification of the biotinylated proteins (Fig S4 A-D, Dataset S1) revealed substantial differences in the putative interactomes of astrocytic NL1-3 (Fig 3F, Dataset S1), when normalized to their respective soluble TurboID controls. Similarly, there were substantial differences between proteins proximal to neuronal versus astrocytic NL2 (Fig 3F, Dataset S1).

We bioinformatically compared our data to previously published NL2 interactome data (identified via NL2 co-immunoprecipitation in adult mouse whole brain (Kang et al., 2014), *in vitro* extracellular NL2 APEX proximity proteomics (Loh et al., 2016), and a yeast two-hybrid screen of the NL2 ICD (Poulopoulos et al., 2009)) and found a significant overlap with our datasets for each of the previously published data (Fig 4A). We also found noticeable differences between the astrocytic and neuronal interactome of NL2. For example, neuronal NL2 is well characterized for interacting with the inhibitory synapse scaffold protein, Gephyrin (Poulopoulos et al., 2009), which we detected as highly enriched in the neuronal NL2 iBioID interactome (Fig 4B, Dataset S1). However, even though Gepyhrin is present in astrocytes at both the mRNA (Zhang et al., 2014) and protein levels (Dataset S1), we did not detect a significant enrichment of Gepyhrin in the astrocytic NL2 iBioID (Fig 4B, Dataset S1). This result strongly suggests cell-type-specific regulation of protein-protein interactions of NL2 and is consistent with our finding that Gephyrin interaction with NL2 is not necessary for astrocyte morphogenesis (Fig 2A, B).

**Figure 4:**
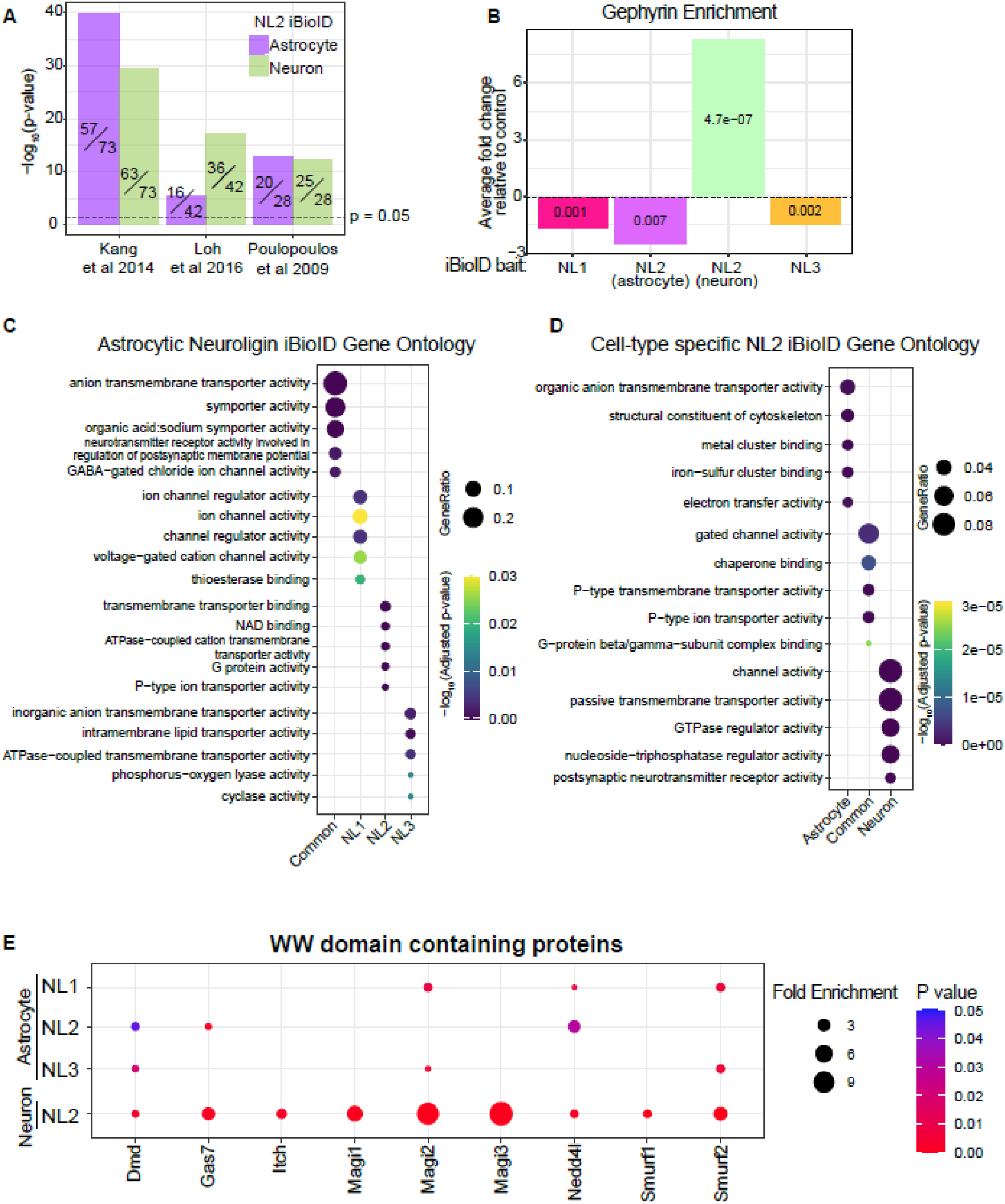
Neuroligin iBioID reveals novel intracellular domain interacting proteins. A) Enrichment of NL2 iBioID datasets with previously published datasets of NL2-interacting proteins. P-values are the result of Fisher’s exact test. The number on the bars indicates the number of proteins from appropriate NL2 iBioID that are found in the published dataset. B) Gephyrin enrichment with each NL iBioID. Enrichment is defined as fold change bait / cytosolic control (Turbo BirA). P values are the results of t-tests within each experiment. C) Top 5 Gene Ontology Biological Processes categories for each NL BioID. Common indicates proteins found in astrocytic NL1, NL2, and NL3 iBioID. The complete gene ontology list can be found in Datset S2. GeneRatio indicates number of gene names (from the input iBioID protein list) in the GO category, out of all gene names in that category. Input for the analysis was proteins from iBioID lists that were at least 1.5 fold enriched at a p-value of <0.05, compared to respective controls. D) Top 5 Gene Ontology Biological Processes categories for each Neuronal or Astrocytic NL2 BioID. Common indicates proteins found in both astrocytic NL2 and neuronal NL2 iBioID. The complete gene ontology list can be found in Dataset S2. GeneRatio indicates number of gene names (from the input iBioID protein list) in the GO category, out of all gene names in that category. Input for the analysis was proteins from iBioID lists that were at least 1.5 fold enriched at a p-value of <0.05, compared to respective controls. E) All identified WW-binding protein interactors in iBioID experiments. Fold change is the enrichment of the interaction with the bait compared to cytosolic Turbo BirA alone, per cell type. P-value represents a two-tailed T-test compared to soluble Turbo BirA, per cell type. Input for the analysis was proteins from iBioID lists that were at least 1.5 fold enriched at a p-value of <0.05, compared to respective controls.

Gene ontology analysis of the unique proximity interactomes across astrocytic NLs suggests distinct functions of NLs in astrocytes (Fig 4C) and of astrocytic vs neuronal NL2 (Fig 4D). These data support our earlier findings, showing that NLs have non-redundant roles in astrocytes and that these differences likely extend beyond the control of astrocyte morphogenesis. Further, gene ontology analysis of NL2-proximal proteins in neurons and astrocytes reveals functional differences. Interestingly, only astrocytic NL2 interacts with proteins that bind actin filaments (Fig 4D, S4E), which are abundant in perisynaptic astrocyte processes (Soto et al., 2023). To facilitate further exploration into the biological mechanisms governing NL control of astrocyte development and function, we have compiled our dataset into a searchable website at https://eroglu-neuroliginbioid.org/.

The putative NL2 interactome we identified enabled us to determine which nearby proteins recognize the PPxY motif in NL2 and thus participate in NL2-dependent astrocyte morphogenesis. To do so, we analyzed WW domain-containing proteins across all datasets that were significantly enriched with the bait protein, compared to the cytoplasmic control. Across the 4 iBioID experiments, we identified 9 WW-domain-containing proteins in NL interactomes (Fig 4E). The WW domain-containing proteins within the NL interactomes varied across NL isoforms and cell types. Even though NL1 and NL2 have identical PPxY motifs (PPDY, Fig 2G), there was minimal overlap in WW domain-containing proteins that interacted with both. This finding suggests that the interactions of NLs with WW domain-containing proteins *in viv*o are not solely determined by the presence of the PPxY motif, and that additional protein-protein interaction motifs in the ICD may confer specificity to these interactions. Furthermore, the utility of including NL3 as a bait in our BioID experiments served as an informative control for the PPxY motif, as NL3 contains the LPxY motif, which is only weakly bound by WW proteins, providing a strong comparative reference for motif-dependent interactions.

### The WW domain-containing E3 ubiquitin ligase Nedd4l is necessary for astrocyte morphogenesis, but not synaptogenesis

The binding of WW domains to WW motifs has various physiological effects in the cell, including cellular growth (Lee et al., 2020), endocytosis of membrane proteins (Totland et al., 2017), and transcription (Sudol et al., 2001). Importantly, WW domain-containing proteins and their interactions with WW motifs have been implicated in various diseases, including the neurodevelopmental disorder Rett Syndrome (Rodrigues et al., 2020), Muscular Dystrophy (Ilsley et al., 2002), and hypertension (Minegishi et al., 2016). We observed three significantly enriched WW domain-containing protein interactors of astrocytic NL2, Dystrophin (*Dmd*), growth arrest-specific protein 7 (*Gas7*), and neural precursor cell expressed developmentally downregulated 4-like protein (*Nedd4l*, also called Nedd4-2) (Fig 4E). We chose to focus on Nedd4l for two reasons. First, both NL1 and NL2 PPDY motifs are necessary for astrocyte morphogenesis (Fig 2I-J), and Nedd4l is enriched in astrocytic NL1 and NL2, but not NL3 (Fig 4E) proximity proteomes. Second, Nedd4l has a known role in regulating astrocyte membrane proteins. Nedd4l controls the abundance of the potassium channel Kir4.1, the gap junction protein, Connexin-43 (Altas et al., 2024; Liu et al., 2021) as well as the abundance and surface localization of the glutamate transporter Glt-1 (Zhang et al., 2017), via ubiquitination.

To investigate if Nedd4l is necessary for astrocyte morphogenesis, we designed and validated an shRNA (Figs 5A, S5A-B) targeting *Nedd4l*, and expressed it in astrocytes cultured with wild-type neurons. We observed a striking reduction in astrocyte morphological complexity in shNedd4l-transfected astrocytes, compared to control (Fig 5A-B). Co-knockdown of *Nedd4l* and *Nlgn2* (using a plasmid that also encodes mCherry) did not further reduce astrocyte complexity. These findings suggest that Nedd4l and NL2 might work in the same pathway (Fig 5B).

**Figure 5:**
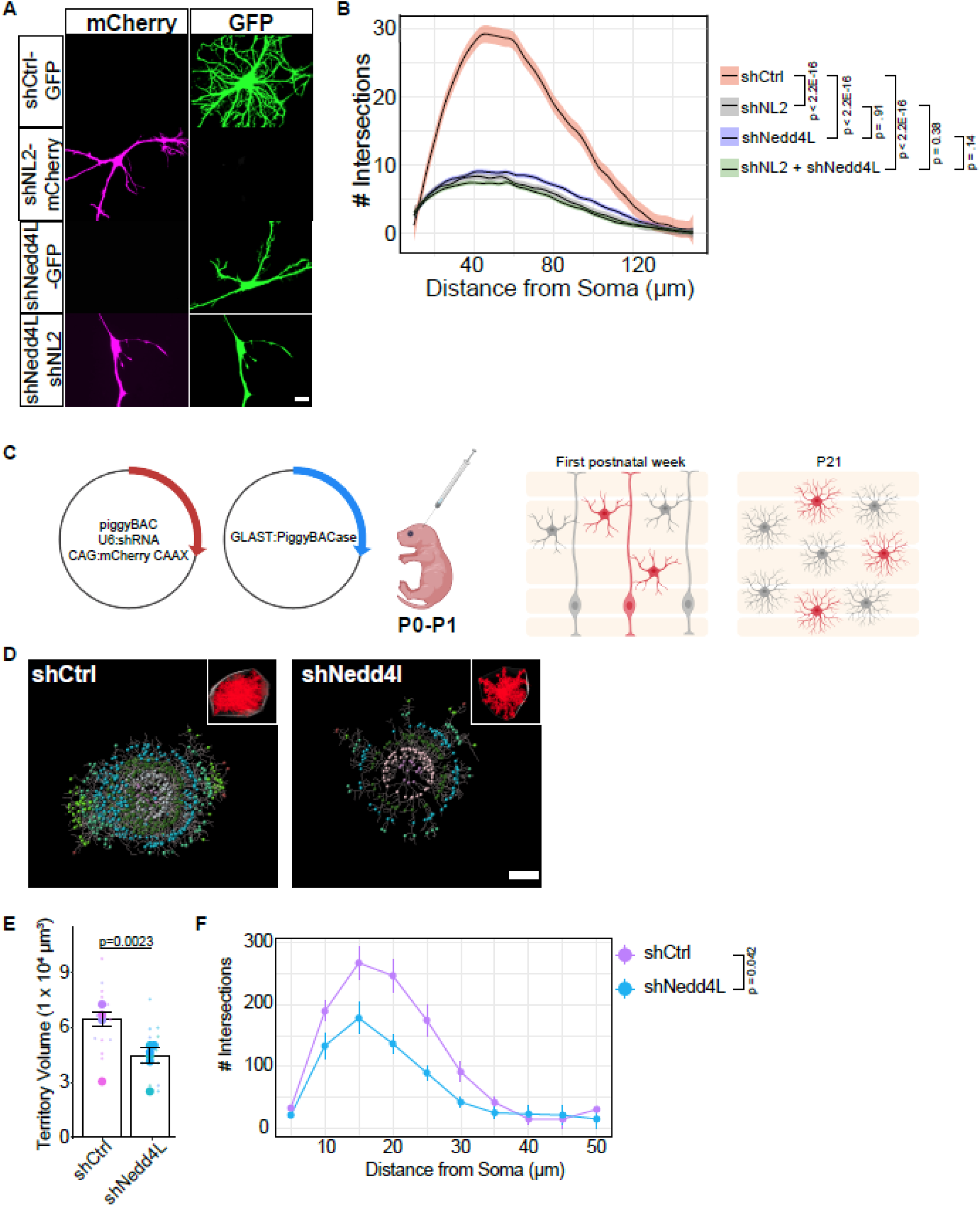
WW domain-containing proteins are necessary for astrocyte morphogenesis in vitro. A) Representative images of astrocytes transfected with either a control shRNA (scrambled NL2 shRNA), an shRNA targeting Nedd4l (also expressing GFP) or shNL2 (also expressing mCherry), or in combination. Astrocytes are cultured on top of neurons (not labeled). Scale bar is 20µm. B) Sholl analysis of the conditions in A. Linear-mixed model ANOVA reveals the main effect of condition F(3, 332)=217.36, p<2.2E-16. P-values on the graph represent Tukey’s posthoc comparison to shCtrl. N = 60-100 cells per condition, across 4 independent culture experiments. Shaded area indicates standard error. C) Cartoon of postnatal astrocyte labeling by electroporation (PALE). Two plasmids (U6:shRNA and CAG:mCherry CAAX flanked by piggyBAC transposons, and GLAST: piggyBACase transposase) are electroporated into late P0 or early P1 mouse lateral ventricles. Brains are collected at P21 and stained for mCherry to visualize astrocyte territories. D) Imaris reconstructions of *in vivo* astrocyte territory volumes (inset) and 3D Sholl analysis of mouse cortical astrocytes. ‘Filament’ cytoskeletal structure is shown with grey lines, while spheres indicate 3D Sholl concentric circles [different colors indicate different distances from the nucleus]. Scale bars = 10µm. E) Territory volume super plot quantification of the conditions in C. Large dots on graph = average per animal. Smaller dots = individual cells. Data were analyzed using a nested ANOVA; main effect of condition, F(1, 8) = 11.95, p=0.0023. The same cells were measured for territory volume and Sholl analysis. N=14-18 cells across 4-7 animals (biological replicates) per condition. Layer identity of cells imaged: shCtrl [L2/3: 1, L4: 10, L5/6: 7]; shNedd4l [L2/3: 2, L4: 8, L5/6: 4] F) 3D Sholl analysis. Linear mixed model ANOVA F(1,9)=5.59, p=0.0424. The same cells were measured for territory volume and Sholl analysis. N=14-18 cells across 4-7 animals (biological replicates) per condition. Error bars indicate standard error.

Next, we tested whether Nedd4l is necessary for astrocyte morphogenesis *in vivo*. To achieve genetically targeted and sparsely labeled astrocytes in the mouse cortex, we used postnatal astrocyte labeling by electroporation (PALE) (Baldwin et al., 2021; Stogsdill et al., 2017). In this method, plasmid DNA expressing a shRNA with CAG:mCherry-CAAX and a GLAST promoter-driven piggyBac enzyme are injected into P0-P1 mouse lateral ventricles and then electroporated into the cortex. This technique results in the uptake of the plasmid by radial glia and subsequent astrocyte labeling after differentiation (Fig 5C). We previously found that reducing astrocytic NL2 by PALE, either by using an shRNA or a floxed allele for *Nlgn2* with astrocytic Cre expression, results in a significantly stunted astrocyte territory volume and branch complexity (Stogsdill et al., 2017). We used confocal imaging of the entire astrocyte in 100µm tissue sections and reconstructed the astrocyte volume using Imaris software. Quantification of the astrocyte territory volume revealed a significant reduction in shNedd4l astrocytes, compared to a scrambled shRNA control (shCtrl) (Fig 5D-E). Additionally, quantification of the astrocyte complexity using *in vivo* Sholl analysis revealed a significant decrease in complexity of Nedd4l knockdown astrocytes (Fig 5F). These data suggest that astrocyte morphogenesis is controlled by Nedd4l at both the level of territory size and branch complexity.

We previously showed that loss of *Nlgn2* in single astrocytes in the mouse V1 cortex also results in reduced excitatory synaptogenesis in the surrounding neuropil. Thus, to test if Nedd4l loss also affects synapse density, we used PALE to knock down *Nedd4l* in sparse astrocytes in the mouse V1 cortex. At P21, we stained cortical sections for presynaptic Bassoon, and postsynaptic PSD95 and Gephyrin. With this strategy, we quantified both excitatory (closely apposed Bassoon and PSD95) and inhibitory (Bassoon and Gephyrin) synapses in the same sections (Fig S5C). We used the mCherry signal from the piggyBAC vector (Fig 5C) to define astrocyte territories. Using open-source SynBot software (Savage et al., 2024), we quantified the synaptic density within and outside of labeled astrocytes (Fig S5C-D). These data revealed no significant differences in synapse density in *Nedd4l*-knockdown astrocytes, *in vivo* (Fig S5E-F).

This data suggests Nedd4l and its interaction with NL2 controls astrocyte morphological development independently from the role of astrocytic NL2 in synaptogenesis, and Nedd4l does not have an independent role in astrocyte-dependent synaptogenesis.

### The PPDY motif in Neuroligin-2 is sufficient for Nedd4l interaction

Nedd4 proteins exhibit preferential binding to PPxY motifs and only weakly bind to LPxY (Hatstat et al., 2021). In the absence of a PPxY containing substrate, Nedd4l is known to bind its own LPxY motif, autoubiquitinate and degrade (Bruce et al., 2008). Because NL1 and NL2 have a PPxY motif, and NL3 has an LPxY motif (Fig 2G; Fig S2B), we wondered if Nedd4l preferentially and directly interacted with NL1 and NL2 ICDs but not the NL3 ICD. To test this, we used either the wildtype NL2 or NL2 ECD/NL1 ICD chimera (Fig 2D) or the mutated PPDY motif constructs that abolish the WW domain-containing motif, as described earlier (Fig 2H-J; Fig 6A). Conversely, we also mutated the NL3 ICD LPDY motif to PPDY to mimic NL1 and NL2 ICDs. All chimeric constructs contained the NL2 ECD to rule out the effects of sequence divergence in NL ECDs.

**Figure 6:**
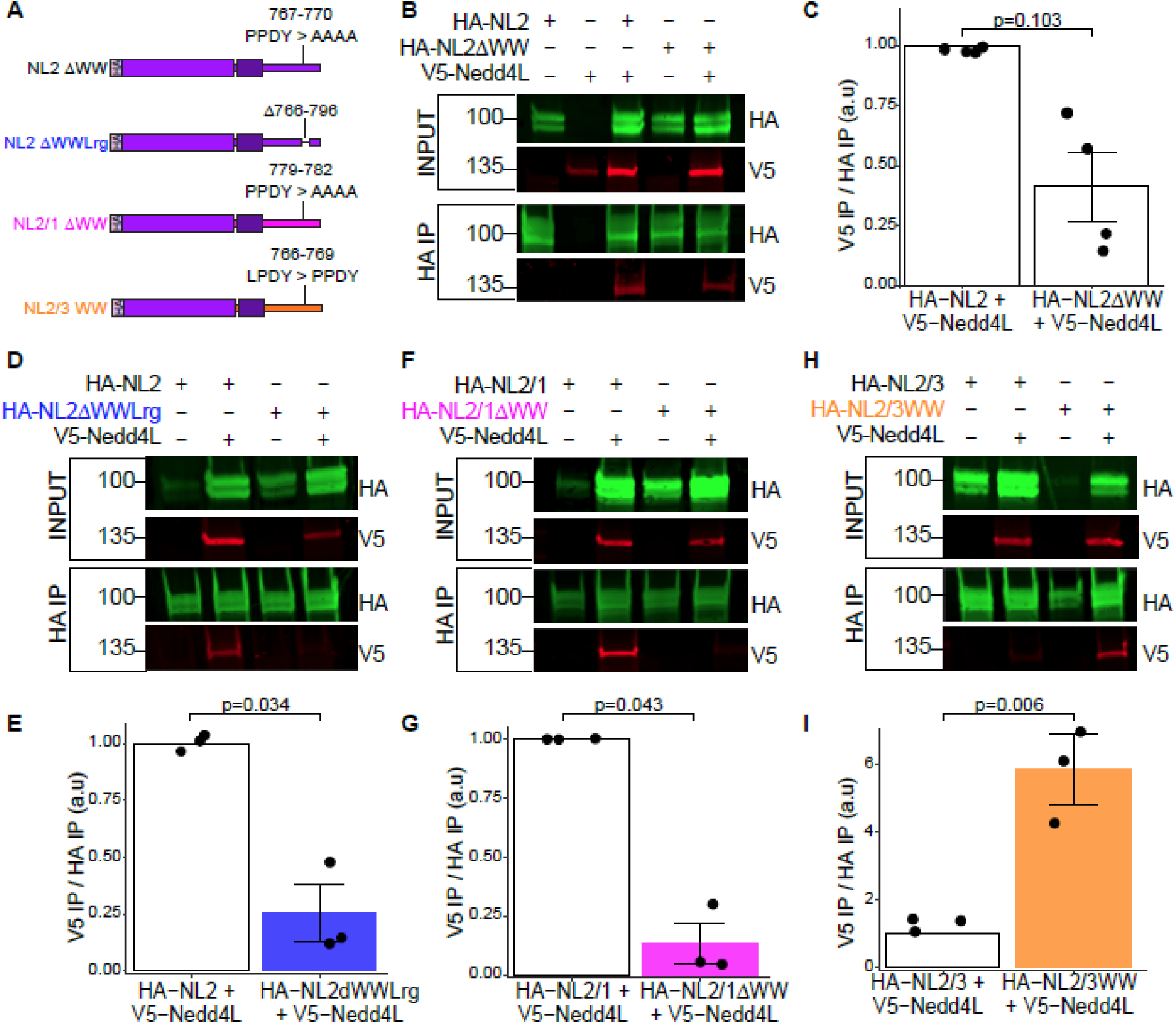
Nedd4l binding to Neuroligins is modulated by intracellular PPxY motifs A) Cartoon of NL2 and NL2-ICD chimeric constructs. Numbers indicate amino acid positions in mouse sequence. B) Western blots from HEK cell lysates transfected with HA-NL2 or HA-NL2ΔWW with or without V5-Nedd4l constructs. Co-immunoprecipitation (IP) of V5-Nedd4l was detected with NL2-HA constructs using Li-Cor fluorescent imaging. C) Densitometry quantification of IP signal from conditions in B. N=4 biological replicates. P-values represent results of paired, Student’s t-test. D) Western blots from HEK cell lysates transfected with HA-NL2 or HA-NL2ΔWWLrg (Deletion of amino acids 766-796) with or without V5-Nedd4l constructs. Co-immunoprecipitation (IP) of V5-Nedd4l was detected with NL2-HA constructs using Li-Cor fluorescent imaging. E) Densitometry quantification of IP signal from conditions in D. N=3 biological replicates (independent culture experiments). P-values represent results of paired, Student’s t-test. F) Western blots from HEK cell lysates transfected with HA-NL2/1 or HA-NL2/1ΔWW with or without V5-Nedd4l constructs. G) Densitometry quantification of IP signal from conditions in F. N=3 biological replicates. P-values represent results of paired, Student’s t-test. H) Western blots from HEK cell lysates transfected with HA-NL2/3 or HA-NL2/3-WW with or without V5-Nedd4l constructs. I) Densitometry quantification of IP signal from conditions in H. N=3 biological replicates. P-values represent results of paired, Student’s t-test. (B, D, F, H) All western blots were run at least 3 times with independent samples. Note that Nedd4l (V5) blots were run separately (eluate divided evenly) to avoid bleed-through from the strong HA signal.

Next, we co-expressed HA-tagged NL2 or NL2 chimeras with V5-tagged Nedd4l in HEK 293T cells and co-immunoprecipitated NLs using anti-HA tag magnetic beads. We found that Nedd4l co-immunoprecipitated with wildtype NL2 (Fig 6B); however, ablation of the PPDY motif in NL2 ICD (NL2ΔWW, Fig 6A) resulted in a substantial, yet variable, reduction of Nedd4l co-immunoprecipitation (Fig 6B, C). A previous study investigating the interaction of NL2 and the WW domain-containing protein S-SCAM (*Magi2*) found that a longer stretch of amino acids (766-796), which contains the PPDY motif, in the NL2 ICD is necessary for direct binding of S-SCAM to NL2 (Sumita et al., 2007). Thus, we next deleted these amino acids from the NL2 ICD (NL2ΔWWLrg, Fig 6A) and performed co-immunoprecipitation with Nedd4l. Loss of this sequence in NL2 ICD completely abolished the interaction with Nedd4l (Fig 6D-E), demonstrating that this is a critical region in NL2 ICD containing the PPDY motif that mediates Nedd4l binding.

Ablation of the PPDY motif in NL1 ICD (NL2/1ΔWW, Fig 6A), however, abolished the interaction with Nedd4l entirely, further showing the importance of the PPDY motif in NL-Nedd4l interactions (Fig 6F-G). As expected, we found that Nedd4l does not interact with wild-type NL3 ICD (Fig 6H). However, mutating the LPDY motif of NL3 ICD to PPDY (NL2/3WW, Fig 6A, H-I) was sufficient to induce an interaction between Nedd4l and NL3 ICD. These results reveal that the PPDY is sufficient for Nedd4l interactions with NL ICDs. Additional sequences in the NL2 ICD between amino acids 766-796 flanking the PPDY motif are also involved in NL2/Nedd4l binding.

### Nedd4l ubiquitinates Neuroligin-2 at K749, resulting in increased stability of Neuroligin-2

Nedd4l is an E3 ubiquitin ligase of the HECT domain family, which has previously been shown to polyubiquitinate the astrocyte membrane proteins Kir4.1, Connexin-43, and Glt-1, resulting in their degradation (Altas et al., 2024; Liu et al., 2021; Zhang et al., 2017). Previous proteomic studies have identified that the intracellular domain of NL2 is ubiquitinated at lysine 749 (K749) (Na et al., 2012; Wagner et al., 2012) in both mouse and rat brain tissues. However, the function of this post-translational modification in NL2 is unknown. Thus, we investigated whether NL2 is ubiquitinated at K749. To do so, we first blocked proteasomal degradation using the proteasomal inhibitor MG-132. We next heat-denatured lysates to disrupt protein-protein interactions and subsequently immunoprecipitated HA-NL2 from primary rat astrocytes. After running the immunoprecipitated eluates on a western blot, we found that HA-NL2 showed a smear of high molecular weight HA-immunoreactive protein, indicative of polyubiquitinated protein (Fig 7A). We found that we could significantly decrease this smear by mutating K749 to R (Arginine), as immunoprecipitation of HA-NL2K749R only showed one predominant band at the molecular weight of NL2, indicating this smear is likely ubiquitination (Fig 7A-B).

**Figure 7:**
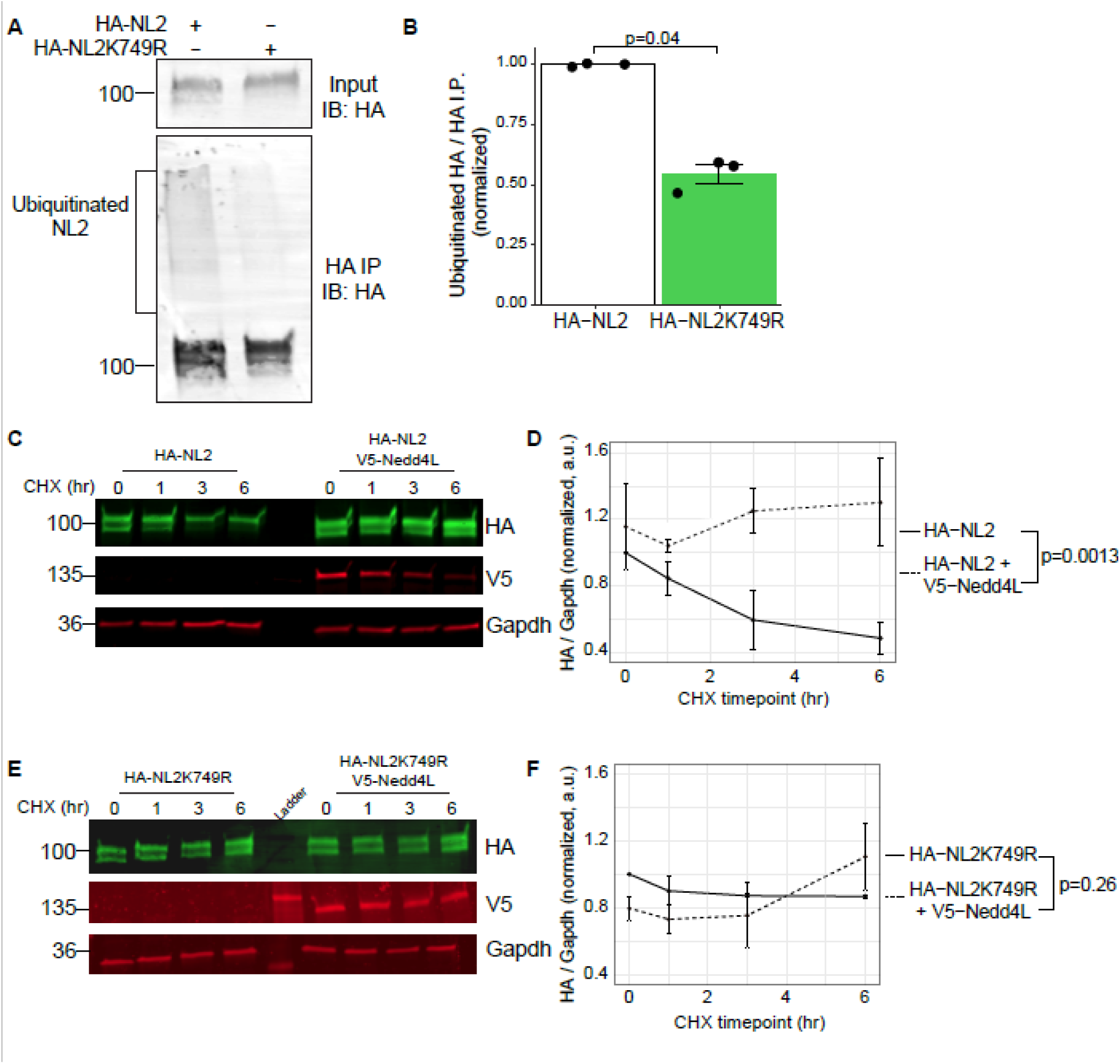
Nedd4l stabilizes Neuroligin-2 via ubiquitination at K749R. A) Western blot from rat primary astrocyte cell lysates and HA-NL2 IPs shows NL2 is ubiquitinated in astrocytes, which is lost when NL2 K749 is mutated to arginine (K749R). Input is 2% of total lysate. B) Quantification of HA ubiquitination signal from conditions in A. The ubiquitinated HA bands >100kDa were quantified via densitometry and normalized to the HA IP band (∼100kDa) per biological replicate. N=3 biological replicates (independent culture experiments). P-values represent results of paired, Student’s t-test. C) Western blots from HEK cell lysates transfected with HA-NL2 or HA-NL2 + V5-Nedd4l and treated with CHX. Numbers above the blot indicate hours after CHX treatment. D) Quantification of the conditions in B. Samples were collected on N=3 independent samples. Two-way ANOVA reveals main effect of condition F(1, 16)=15.018, p=0.00134, with no significant effect of time-point F(3, 16)=0.41, p=0.74827 or interaction F(3, 16)=2.242, p=0.123. E) Western blots from HEK cell lysates transfected with HA-NL2K749R or HA-NL2K749R + V5-Nedd4l and treated with CHX. Numbers above the blot indicate hours after CHX treatment. F) Quantification of the conditions in D. Samples were collected on N=3 independent samples. Two-way ANOVA reveals no significant effect of condition F(1, 16)=1.392, p=0.255, time-point F(3, 16)=1.681, p=0.211 or interaction F(3, 16)=2.328, p=0.113.

We next aimed to test if Nedd4l ubiquitinates NL2. Interestingly, while the *Nedd4l* expression construct works well in HEK 293T cells, we were unable to overexpress Nedd4l in primary astrocytes. This may be caused by global proteasome disruptions and cytotoxicity due to Nedd4l’s role in astrocyte protein turnover (Altas et al., 2024). Therefore, we used HEK 293T cells to express HA-NL2 with or without V5-Nedd4l and probed for GFP-tagged ubiquitination by western blot using a GFP antibody. We found that, compared to NL2 alone, co-expression with Nedd4l resulted in a high-molecular-weight smear above the expected NL2 size, indicating that Nedd4l is sufficient to enhance polyubiquitination of NL2 (Fig S5G-H). Further, we could reduce ubiquitination by mutating NL2 K749 to arginine (K749R). Co-expression of Nedd4l and NL2K749R did not result in strongly enhanced polyubiquitination of NL2, indicating that K749 is the major ubiquitination residue in NL2, but others may be present (Fig S5G-H). Together, these data reveal that Nedd4l is sufficient to ubiquitinate NL2 at K749.

While ubiquitination is typically associated with protein degradation, it can also serve as a post-translational signal for various cellular processes (Mukhopadhyay and Riezman, 2007). We first tested whether Nedd4l-mediated ubiquitination of NL2 leads to its degradation. We expressed NL2 with or without Nedd4l in HEK cells and collected cell lysates at different time points after cycloheximide (CHX) treatment, which blocks nascent translation. Western blotting of these samples revealed a decrease in HA-NL2 abundance over time, indicating NL2 is degraded (Fig 7C-D). Strikingly, co-expression of Nedd4l with NL2 completely blocked this degradation and stabilized the NL2 protein over time (Fig 7C-D). Interestingly, the NL2K749R ubiquitination mutant was sufficient to block NL2 degradation, with no further effect by the addition of Nedd4l (Fig 7E-F).

Taken together, these data reveal that ubiquitination at K749 controls NL2 degradation; however, ubiquitination at K749 by Nedd4l stabilizes the NL2 protein and prevents its degradation.

### Ubiquitination of Neuroligin-2 at Lysine 749 is required for astrocyte morphogenesis

NL2 is ubiquitinated *in vivo* (Na et al., 2012; Wagner et al., 2012), yet whether ubiquitination at this residue has a functional role is unexplored. Therefore, we next tested if ubiquitination at K749 is necessary for NL2’s control of astrocyte morphogenesis. To do so, we expressed NL2K749R (also mutated to be resistant to shNL2) in primary astrocytes and subsequently co-cultured the astrocytes on top of neurons to quantify astrocyte complexity. Surprisingly, we found that NL2K749R expression significantly reduced astrocyte morphology in a dominant-negative fashion compared to the shRNA control (Fig 8A-B). NL2K749R was incapable of any morphological rescue of shNL2-transfected astrocytes (Fig 8A-B).

**Figure 8:**
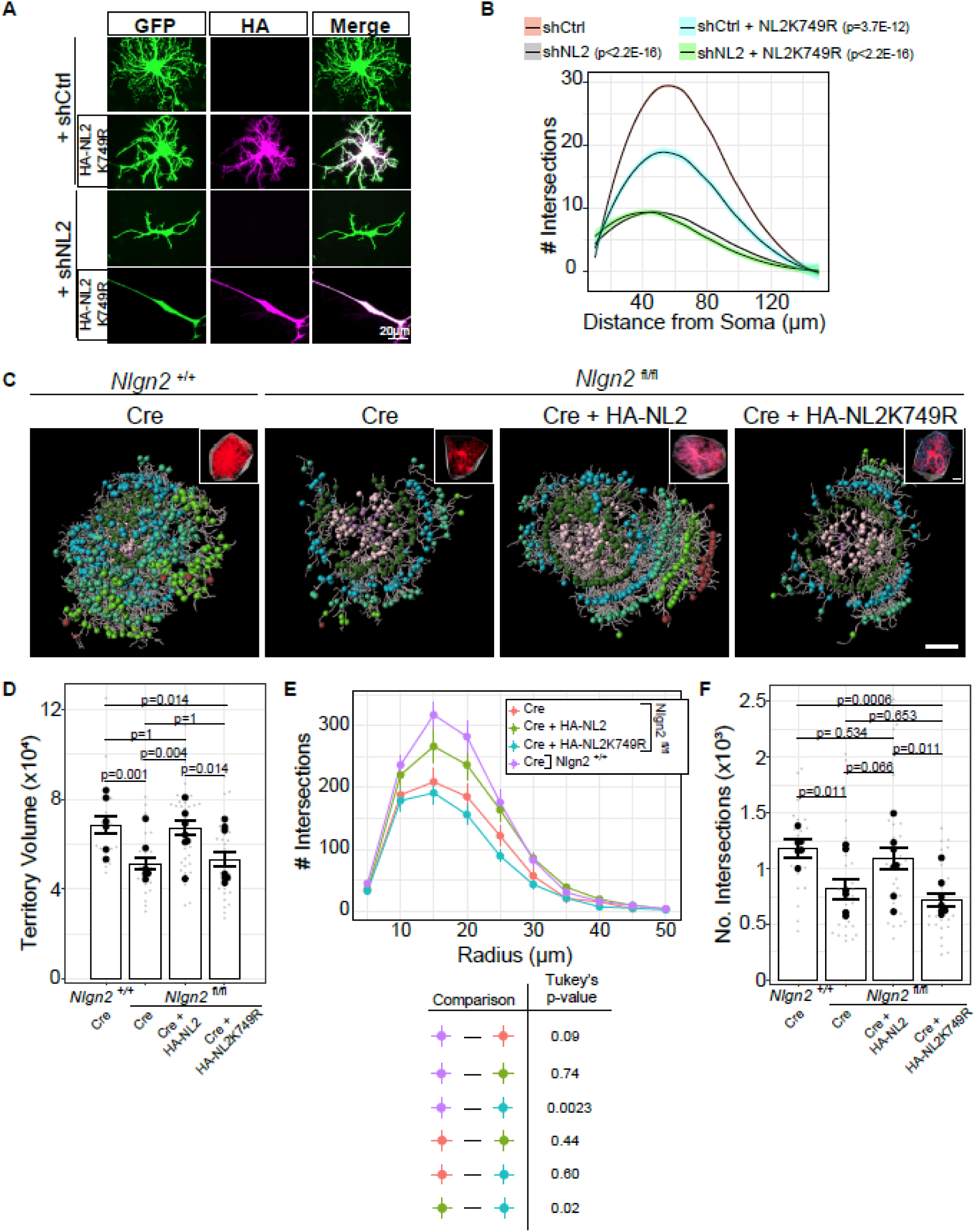
Neuroligin-2 ubiquitination at K749 is critical for astrocyte morphogenesis. A) Representative images of astrocytes transfected with either shCtrl or shNL2 with or without HA-NL2K749R expression. Scale bars are 20µm. B) Sholl analysis quantification of images in J. Linear-mixed model ANOVA reveals the main effect of condition F(3, 284)=103.14, p<2.2E-16. P-values represent Dunnett’s posthoc with respect to shCtrl. N = 62-80 cells per condition, across 3 independent culture experiments. Shaded area indicates standard error. C) Representative images of reconstructed astrocytes. ‘Filament’ cytoskeletal structure is shown with grey lines, while spheres indicate 3D Sholl concentric circles [different colors indicate different distances from the nucleus]. Insets show the astrocyte signal [tdTomato signal in red; HA signal in cyan] with the convex hull [grey surface lines]. Scale bar = 10µm. D) Territory volume super plot quantification of the conditions in C. Large black dots on graph = average per animal. Smaller grey dots = individual cells. Data were analyzed using Kruskal-Wallis rank sum test: χ^2^ = 19.5, df=3, p=0.00013. P-values on the graph are results of the Wilcoxon rank sum post hoc tests with Holm-Šídák multiple comparison correction. The same cells were measured for territory volume and Sholl analysis. N=21-33 cells across 6-7 animals (biological replicates) per condition. E) 3D Sholl analysis. Linear mixed model ANOVA F(3, 102)=5.08, p=0.0025. P-values in the table represent Tukey’s post hoc tests. The same cells were measured for territory volume and Sholl analysis. N=21-33 cells across 6-7 animals (biological replicates) per condition. Error bars represent standard error. F) Total number of intersections (No. Intersections, y-axis) from cells in Sholl Analysis (E). Large black dots on graph = average per animal. Smaller grey dots = individual cells. Data were analyzed using Kruskal-Wallis rank sum test: χ^2^ = 19.247, df=3, p=0.000243. P-values on the graph are results of the Wilcoxon rank sum post hoc tests with Holm-Šídák multiple comparison correction. The same cells were measured for territory volume and Sholl analysis. N=21-33 cells across 6-7 animals (biological replicates) per condition.

Finally, we investigated whether NL2K749R is crucial for astrocyte morphogenesis in the developing mouse cortex. In *Nlgn2* homozygous flox animals (*Nlgn2* ^fl/fl^), also carrying a Rosa26:lox-STOP-lox tdTomato allele, we used PALE (Fig 5C) to express Cre to ablate *Nlgn2* in astrocytes, while simultaneously expressing HA-NL2 (WT) or HA-NL2K749R (Fig 8C). At P21, we harvested the brains to analyze astrocyte morphological development via Imaris reconstruction and 3D Sholl analysis. As previously shown (Stogsdill et al., 2017), the knock-out of both copies of *Nlgn2* via Cre expression in astrocytes results in a significantly diminished territory volume, compared to *Nlgn2* ^+/+^ animals (also carrying the Rosa26:lox-STOP-lox tdTomato allele) (Fig 8D).

This morphological deficit was rescued by co-expressing an HA-tagged NL2 plasmid in the astrocytes (Fig 8D). Supporting our *in vitro* findings (Fig 8B), co-expression of HA-NL2K749R with Cre in *Nlgn2* ^fl/fl^ astrocytes was not able to rescue the reduction in territory volume and showed no significant difference from *Nlgn2* ^fl/fl^ astrocytes that received only Cre (Fig 8D). We observed a bimodal distribution in the territory volumes of *Nlgn2* ^fl/fl^ astrocytes that received Cre + HA-NL2K749R, which was not related to sex or astrocyte layer identity, suggesting that an unknown variable influences the functional importance of NL2 ubiquitination status.

Using 3D Sholl analysis (Fig. 8E), we observed a trend toward increased branch complexity in HA-NL2–expressing astrocytes compared to Nlgn2 ^fl/fl^ astrocytes expressing Cre alone; however, this difference did not reach statistical significance. In contrast, astrocytes expressing HA-NL2K749R showed no improvement in branch complexity relative to *Nlgn2*-deficient astrocytes. We next calculated the total number of intersections as a summary measure of astrocyte complexity (Fig. 8F). This analysis revealed that *Nlgn2* deletion significantly reduces total branch complexity, which is restored by HA-NL2 but not HA-NL2K749R expression. Notably, HA-NL2K749R-expressing wildtype astrocytes also remained significantly less complex than both wild-type controls and HA-NL2 rescue astrocytes.

Together with the robust rescue of territory volume (Fig. 8D), these findings demonstrate that wild-type NL2 restores astrocyte morphogenesis, but ubiquitination-deficient NL2K749R mutant fails to do so. These results support a model in which Nedd4L-mediated ubiquitination of NL2 is required for its proper function in regulating astrocyte morphogenesis *in vivo*.

## DISCUSSION

Astrocytes play crucial roles in nervous system development and function. These functions rely on astrocytes’ highly branched morphology, which is largely controlled by cell-adhesion molecules (Ackerman et al., 2021; Baldwin et al., 2021; Stogsdill et al., 2017; Tan et al., 2023). However, the mechanisms downstream of these cell adhesion molecules are not fully understood. Here, we show that NLs are non-redundant in controlling astrocyte morphology due to their unique intracellular protein interactomes. We found a critical motif that regulates Nedd4l interaction and ubiquitination of NL2, which is necessary for NL2-mediated control of astrocyte morphogenesis. Further, we provide *in vivo* astrocytic NL1-3 and neuronal NL2 proximity proteomes, which provide insights into the differences in astrocytic NL function and cell-type-specific functions of NL2 between astrocytes and neurons.

How do NLs uniquely contribute to astrocyte morphogenesis? We found that overexpression of NLs could not compensate for the loss of one another *in vitro*, which may be due to several factors. First, NL mRNA expression patterns differ in the developing cortex (Clarke et al., 2018), suggesting that temporal dynamics of NLs may differentially control astrocyte cytoarchitecture. Second, NLs form both homo– and heterodimers, and dimerization is necessary for both localization and function (Poulopoulos et al., 2012; Shipman and Nicoll, 2012). In neurons, only NL1/NL3 and NL1/NL2 heterodimers have been detected (Poulopoulos et al., 2012); however, our results indicate that both NL1/NL2 and NL2/NL3 heterodimers are present in both astrocytes and neurons, suggesting that cell-type-specific heterodimers may impact NL function during brain development. Lastly, our interactome data reveal distinct protein interactions of NLs, which likely contribute to unique astrocytic NL function. Interestingly, gene ontology analysis revealed a significant interaction with actin-binding proteins and cytoskeletal-related proteins specifically in astrocyte NL2, and minimally in astrocyte NL1, NL3, or neuronal NL2, suggesting that astrocytic NL2 plays a dominant role in controlling cell morphology (Dataset S2, Fig S4 E-F). It is important to note a few limitations of our iBioID data: first, because the TurboID promiscuously biotinylates proteins within 10nm, interactions with a bait protein (here, NLs) may or may not be due to direct binding and should be validated orthogonally, as we showed with Nedd4l (Fig 6). Second, interactions with NLs may be transient and/or developmentally regulated, and thus, our 3-day biotinylation treatment window may capture both static and transient interactions. Therefore, these data should not be interpreted as a compendium of NL-interacting proteins. Lastly, GFAP promoter activity is highest in the lower layers of the cortex (Fig 3C-D), thus our astrocyte-specific proteomic experiments are largely influenced by lower-layer astrocytes. Nevertheless, the iBioID interactomes generated in this study will be beneficial in elucidating the mechanisms through which NLs control astrocyte structure and function.

In this work, we show that the WW domain-containing protein Nedd4l is critical for astrocyte morphogenesis, and this protein has distinct specificity for each NL, which we validated by coimmunoprecipitation. How do WW domain proteins specify interactions with NLs? Nedd4l preferentially binds PPxY motifs, but can also weakly bind LPxY (Bruce et al., 2008; Hatstat et al., 2021). We identified that NL1 and NL2 contain a PPxY motif in their ICDs, whereas NL3 contains an LPxY motif. This single amino acid change ultimately affects Nedd4l’s binding to NLs but does not fully explain the interaction specificity. Only after deleting amino acids 766-796 in NL2 is Nedd4l binding blocked, suggesting a larger surface is critical for the interaction between NL2 and Nedd4l. This may explain why mutation of PPDY to AAAA in the NL2 ICD still restores some astrocyte complexity (Fig 2J) and suggests that the WW-binding motif in NL2 is not the sole determinant of NL2 function in astrocyte morphogenesis. Moreover, in our iBioID results, Nedd4l is not enriched in the NL1 proximal proteome as strongly as NL2 in astrocytes, which may indicate a weaker physiological interaction. Tyrosine phosphorylation in WW motifs is known to occlude binding with WW domains (Chen et al., 1997). The conserved Tyrosine (Y) in the PPxY motif is the same residue that modulates Gephyrin binding (Y770) (Poulopoulos et al., 2009) and is phosphorylated in neuronal NL1 but not neuronal NL2 (Letellier et al., 2018), therefore, phosphorylation and/or Gephyrin binding may modulate Nedd4l binding to NLs. Nedd4l interactions with its known targets are controlled through adapter proteins, namely the 14-3-3 family (Lee et al., 2018; Nagaki et al., 2006). Our data reveal that astrocytic NL2 interacts with several members of the 14-3-3 family, which may influence Nedd4l-NL interactions in astrocytes. Lastly, several WW domain-containing proteins interact with NLs in a cell-type-specific manner (Fig. 4E). Future studies will reveal the cell-type-specific effects of WW domain-containing protein interactions with NLs.

Until this study, the functional consequence of NL2 ubiquitination had not been demonstrated in either astrocytes or neurons. We found that Nedd4l ubiquitinates NL2 at K749. Numerous reports have shown that Nedd4l ubiquitinates the astrocytic targets Kir4.1 and Connexin-43 (Altas et al., 2024), resulting in proteasome-mediated degradation (Liu et al., 2021; Zhang et al., 2017; Zhou et al., 2007). However, here we found that Nedd4l ubiquitination of NL2 stabilizes NL2. Interestingly, ubiquitination-mediated protein stabilization has been reported previously for other HECT E3 ubiquitin ligases (Xu et al., 2015). We speculate that Nedd4l-mediated NL2 stability may maintain the surface pool of NL2 and/or overall NL2 abundance, thereby supporting synaptic contacts with Neurexins.

While Nedd4l ubiquitination of NL2 does not lead to protein degradation, our data reveal that NL2K749 ubiquitination can be a degradative signal. In HEK cells, NL2 protein abundance significantly decreases over time, and blocking ubiquitination at K749 prevents this degradation (Fig 7C-F). The functional consequences of ubiquitination at NL2 may differ based on the E3 ubiquitin ligase that targets NL2 and/or the subcellular localization of the ubiquitination event. It is possible that NL2 is ubiquitinated by Nedd4l at the cell membrane, where Nedd4l is localized, leading to NL2 stabilization. If Nedd4l cannot ubiquitinate NL2, NL2 may be internalized and targeted for degradation by another E3 ubiquitin ligase. It will be important to disentangle the type of ubiquitin chain attached to NL2 in the presence or absence of Nedd4l.

A limitation of this experiment is that it was performed in heterologous cells, due to the low expression of astrocytic NL2 in the absence of neurons, possibly resulting from rapid degradation. Future work using cell-type-specific knock-in or conditional degradation models will be required to test Nedd4l’s role in regulating NL2 stability in astrocytes, *in vivo*. However, we did detect a high-molecular-weight smear in HA-NL2 in primary rat astrocytes, which is sharply diminished in HA-NL2K749R-expressing cells, highly suggestive of poly-ubiquitination. The identity of this smear can be resolved in the future using direct assays of polyubiquitination, including chain-specific polyubiquitination events. Lastly, these findings also provide insight into how perturbations in NL2 abundance disrupt astrocyte morphology. We find that the NL2K749R mutation (Fig 8A-B), which may stabilize the NL2 protein in astrocytes by preventing proteasome-mediated degradation, acts in a dominant-negative manner (Fig 7C-D). Interestingly, we do not observe an effect on astrocyte morphology with overexpression of wild-type NL2 (Fig S1C, D), suggesting that excessive NL2 in astrocytes is not sufficient to explain the dominant-negative effect of NL2K749R. Our findings suggest that the balance between NL2 stabilization and degradation is controlled by ubiquitination of K749. We postulate that this modification is crucial for regulating NL2 levels and surface localization to establish proper neuron-astrocyte adhesions and support astrocyte morphogenesis.

Finally, proteomics data revealed that NL2 is ubiquitinated at lysine K749 in both mouse and rat brains, whereas NL1 and NL3 are not (Na et al., 2012; Wagner et al., 2012). We found that blocking ubiquitination at K749 results in a stunted morphological phenotype for NL2-mediated astrocyte morphogenesis, both *in vitro* and *in vivo* (Fig 8A-F). This phenotype may be due to mislocalized NL2, thus leading to improper targeting of perisynaptic astrocyte processes in the neuropil and preventing precise interactions with neuronal neurexins. To this end, understanding how trans-synaptic neurexin binding to NL2 affects Nedd4l interaction with NL2 and/or NL2 ubiquitination is an important future direction. Altogether, this finding underscores the importance of NL2 turnover in the development of astrocyte morphology.

Emerging evidence emphasizes the differential roles of astrocytic versus neuronal NLs. Numerous laboratories have identified physiological roles of NLs in various glia including astrocytes (Stogsdill et al., 2017; Ackerman et al., 2021), oligodendrocyte precursor cells (Li et al., 2024), as well as glioma (Venkatesh et al., 2015, 2017). Recently, a study claimed that astrocytic NLs do not control astrocyte morphogenesis or synapse formation (Golf et al., 2025). Using a tamoxifen-inducible astrocyte Cre line and floxed NL alleles to knock out *Nlgns1-3* in the mouse brain, the authors demonstrate a decrease in astrocyte-specific ribosome-bound NL mRNA only at 8 weeks postnatally but show no difference in cortical astrocyte morphology at P35. We suspect that their incongruous findings are due to incomplete knockdown of NLs early in development, which we have shown to be particularly difficult to knock out *in vivo* using the traditional Cre-Lox system (Stogsdill et al., 2017). In this work, to circumvent such issues, we have used carefully validated shRNAs (Chih et al., 2006) *in vitro* and *in vivo*, which have no compounding effect in NL2 cKO astrocytes, indicating no off-target effects (Stogsdill et al., 2017), and we have validated these findings using the CRISPR-Cas9 system (Fig S1E-H) and PALE-mediated KO and rescue of NL2-dependent astrocyte morphogenesis (Fig 8C-F).

In summary, we provide new insights into NL-mediated control of astrocyte morphogenesis and reveal that the WW domain-containing protein Nedd4l is required for astrocyte morphogenesis by ubiquitinating and stabilizing NL2. Lastly, we show that NL2 ubiquitination is critical for astrocyte morphogenesis, *in vitro and in vivo*.

## Acknowledgments

This work was supported by the National Institutes of Health (R01 NS102237; R01 DA047258 to C. Eroglu, F32 NS112565 to K. Sakers, and F31NS125985 to J.J. Ramirez), and fellowships from The Ruth K. Broad Biomedical Research Foundation (N. Elazar) and the Life Sciences Research Foundation (N. Elazar).

We thank Dr. Roger Nicoll for the NL chimeric constructs. We thank members of Dr. Scott Soderling’s lab for iBioID constructs and protocols. We thank Dr. Dolores Irala for her help with figure organization. We thank Dr. Jessica Moore, Dr. Dolores Irala, Dr. Jeff Stogsdill and Dr. Debra Silver for critical review of this manuscript. C. Eroglu is an HHMI Investigator.

## MATERIALS & METHODS

### Mice

All mice and rats were used in accordance with the Institutional Animal Care and Use Committee (IACUC), under the guidance of the Duke Division of Laboratory Animal Resources (DLAR) with protocol numbers A147-17-06, A117-20-05, and A103-23-04. All animals were housed in 12-hour light/dark cycles. Timed-pregnant CD1 female mice (RRID: IMSR_CRL:022) were used for PALE and iBioID experiments, obtained from Charles River Laboratories. The NL2 floxed (B6;SJL-Nlgn2tm1.1Sud/J; Stock #025544, RRID:IMSR_JAX:025544), and Ai14 (RTM) (B6;129S6-Gt(ROSA)26Sortm14(CAG-tdTomato)Hze/J; Stock #007908, RRID:IMSR_JAX:007908) animals were obtained through Jackson Laboratory. CD Sprague-Dawley IGS Rats (SD-001, RRID: IMSR_CRL:400) were used for primary culture preps, obtained from Charles River Laboratories. Mice and rats of both sexes were pooled and included in all experiments.

### Plasmids

#### shRNA plasmids

shRNAs for NL1(RRID:Addgene_59339), 2 (RRID:Addgene_59358), and 3 (RRID:Addgene_59359) and scrambled NL shRNA control (shCtrl) constructs were described previously(Stogsdill et al., 2017). Antisense shRNA sequences:

Nlgn1: CCCAACACTATACCAGGGATT

Nlgn2: GGAGCAAGTTCAACAGCAA

Nlgn3: GCCCTTGGACTTAACATTTAT

shRNAs were cloned into pLKO.1 vector (a kind gift from David Root, Addgene plasmid #10878, RRID: Addgene_10878). A CAG-eGFP sequence was introduced into the vector for expression in primary astrocytes. Antisense shRNA sequences:

■ Nedd4l: TTTATTAAGGGCTTTGCCTGC (Horizon Discovery, TRCN0000086868)

For PALE, the same shRNA sequence was cloned into a piggyBAC transposon system expressing mCherry CAAX and transfected with a GLAST-piggyBACase plasmid (a kind gift from Dr. Joseph LoTurco) as previously described(Tan et al., 2023). The Nedd4l shRNA in the piggybac backbone is available at Addgene (RRID:Addgene_249589).

#### NL chimeric plasmids

Chimeric HA-NL2 constructs were a kind gift from Dr. Roger Nicoll and described previously(Nguyen et al., 2016) (NL2/1 Chimera RRID_Addgene_249601; NL2/3 Chimera RRID_249602). These constructs were modified to be resistant to shNL2(Stogsdill et al., 2017).

#### HA-NL2 mutation plasmids

All plasmids below were made in the HA-NL2-RM backbone, to be resistant to shNL2.

Deletion of the PDZ domain (RRID:Addgene_249594), poly-proline domain (RRID:Addgene_249595), and mutation of WW domains (PPDY → AAAA in NL2 (RRID:Addgene_249600), Large WW Deletion (a.a. 766-796 deletion, RRID:Addgene_249599), PPDY ◊ AAAA in NL1/2 chimera (RRID:Addgene_249597) or LPDY ◊ PPDY in NL3 chimera (RRID:Addgene_249598)) and K749R (RRID:Addgene_249588) were performed using custom gene blocks (IDT) in the HA-NL2-RM backbone. Mutation of the gephyrin binding residue in HA-NL2-RM (HA-NL2-GBM-RM RRID:Addgene_249596) was performed via site-directed mutagenesis using the QuikChange Lightning kit (Agilent) and the following primers:

Sense: 5’ GCGCCCTGCCTGTgcagccGACgctACCCTGGCCTTGCGCCG 3’

Antisense: 5’ GCCCGGCGCAAGGCCAGGGTAGCGTCggctgcACAGGCAG 3’.

#### TurboID plasmids

pZac2.1-gfaABC1D-Turbo-BirA plasmid was a kind gift from Dr. Scott Soderling. The NL-Turbo fusion constructs were made by PCR amplification from the HA-tagged expression plasmids and subsequent In-Fusion cloning (Takara). A flexible glycine hexamer linker was inserted between the NL expression sequence and the Turbo-BirA. pZac2.1-GFABC1D-Turbo-BirA was digested with XhoI and NotI to release the Turbo-BirA to insert HA-NL1/2/3-Turbo-BirA. The gfaABC1D promoter was swapped for hSyn1 for neuronal expression of NL2-TurboID. The TurboID constructs are available through Addgene: gfaABC1D HA-NL2 Turbo RRID:Addgene_249590; gfaABC1D HA-NL1 Turbo RRID:Addgene_249591; gfaABC1D HA-NL3 Turbo RRID:Addgene_249592; hSyn1 HA-NL2 Turbo RRID:Addgene_249593.

#### Expression plasmids

HA-NL1 (RRID:Addgene_15227), HA-NL2 (RRID:Addgene_15259) and HA-NL3 (RRID:Addgene_59318) were previously described(Stogsdill et al., 2017). V5-Nedd4l was made by custom geneblock cloning (IDT) from the HA-Nedd4l (human sequence, a kind gift from Joan Massague (Addgene plasmid # 27000; RRID:Addgene 27000)). GFP-Ubiquitin was a kind gift from Nico Dantuma (Addgene plasmid # 11928; RRID:Addgene 11928). For NL2 rescue PALE experiments, the CMV promoter was replaced with GfaABC1D(Lee et al., 2008) for both HA-NL2 (RRID:Addgene_249587) or HA-NL2K749R (RRID:Addgene_249588) via In Fusion Snap Assembly (Takara). pCAG-Cre was a gift from Connie Cepko (Addgene plasmid # 13775; http://n2t.net/addgene:13775; RRID:Addgene_13775).

#### CRISPR plasmids

pX601-mCherry was a gift from Yuet Wai Kan (Addgene plasmid # 84039; http://n2t.net/addgene:84039; RRID:Addgene_84039). The CMV promoter was replaced with GfaABC1D promoter from the Turbo-BirA plasmids. SaCas9 gRNAs targeting rat *Nlgn2* were designed using CRISPOR(Concordet and Haeussler, 2018) software, and cloned into the BsaI sites in the pX601 vector.

■ *Nlgn2* gRNA1: AAGAACTGCACGACCGGGCC
■ *Nlgn2* gRNA2: GTGGTGAACACAGCCTACGG (RRID:Addgene_250933)
■ *Nlgn2* gRNA3: GCCACACTCAACTACCGTCT (RRID:Addgene_250934)

For the CRISPR GFP translation assay, we cloned the NL2 gRNAs and their PAM sequences upstream of and in frame with GFP under the control of the chicken beta actin promoter.

### Rat astrocyte and neuron co-culture

Astrocyte and neuron cultures were performed as described(Baldwin et al., 2021; Stogsdill et al., 2017). Briefly, P1 rat cortices were dissected and digested in ∼7.5 units/ml Papain (Worthington). A single-cell suspension was extracted by gentle pipetting with ovomucoid solutions (Worthington). Cells were filtered through a 20µm mesh membrane (Elko Filtering 03-20/14) and then split for astrocyte and neuron cultures.

For neuronal isolation, cortical cells were applied to negative panning dishes coated with Bandeiraea Simplicifolia Lectin 1 (x2), followed by AffiniPure goat anti-mouse IgG+IgM (H+L) (Jackson ImmunoResearch 115-005-044, RRID: AB_2338451), and AffiniPure goat anti-rat IgG+IgM (H+L) (Jackson ImmunoResearch 112-005-044, RRID:AB_2338094) antibodies to bind unwanted cells and debris. Cell suspensions were passed onto positive panning dishes coated with mouse anti-L1 (ASCS4, Developmental Studies Hybridoma Bank, Univ. Iowa, RRID:AB_528349) to bind cortical neurons (>95% neuron). Adherent cells were collected following several washes with DPBS (Gibco 14287) (supplemented with BSA and insulin) and forceful pipetting with a P1000 pipette. Isolated neurons were pelleted (11 minutes at 200 g) and resuspended in serum-free neuron growth media (NGM; Neurobasal, B27 supplement, 2 mM L-Glutamine, 100 U/mL Pen/Strep, 1 mM NaPyruvate, 4.2 μg/mL Forskolin, 50 ng/mL, BDNF, and 10 ng/mL CNTF). 100,000 neurons were plated onto 12 mm glass coverslips (24-well plate) coated with 10 μg/mL poly-D-lysine (PDL, Sigma P6407) and 2 μg/mL laminin, and incubated at 37oC in 10% CO2. On DIV2, neurons were treated with 10μM AraC for one day. Further, on DIV2, media was changed to Neurobasal Plus Growth Media (NGM+; Neurobasal Plus, B27 Plus supplement, 100 U/mL Pen/Strep, 1 mM NaPyruvate, 4.2 μg/mL Forskolin, 50 ng/mL, BDNF, and 10 ng/mL CNTF). Neurons were given a 50% media change every third day until astrocyte co-culture. A detailed protocol can be found at: https://www.protocols.io/view/glia-free-cortical-neuron-culture-36wgq3r35lk5/v1.

For astrocyte cultures, cortical cells were centrifuged and resuspended in astrocyte growth medium (AGM; DMEM (Gibco 11960), 10% FBS, 10 μM, hydrocortisone, 100 U/mL Pen/Strep, 2 mM L-Glutamine, 5 μg/ml Insulin, 1 mM NaPyruvate, 5 μg/ml N-Acetyl-L-cysteine). On the third day in vitro (DIV), astrocytes were washed with DPBS and then shaken vigorously by hand until only adherent astrocytes remained. DPBS was replaced with fresh AGM. AraC was added to the culture at DIV5 for two days. On DIV7, astrocytes were passaged via Trypsin (Gibco) into 6– or 12-well dishes and then transfected on DIV8 with Lipofectamine-LTX. On DIV10, transfected astrocytes were trypsinized and resuspended into NGM+ and plated on neurons at a density of 20,000 cells per well (24 well plates). Cells were fixed after 48 hours of co-culture with warm 4% PFA (Electron Microscopy Sciences) for exactly 7 minutes.

### Immunocytochemistry

After fixation in 4% warmed PFA for 7 minutes (except in the case of surface staining in which cold PFA was used for 5 minutes), coverslips were blocked in a blocking buffer (150 NaCl, 50 Tris-base, 1% BSA, 100 mM L-lysine, 0.04% Sodium azide, pH 7.4) containing 0.2% Triton X-100 (omitted in surface staining) and 50% NGS was added to the coverslips for 30 minutes, followed by 2 washes of DPBS. Primary antibodies were diluted in a blocking buffer containing 10% NGS and added to the coverslips overnight at 4C. The following antibodies were used: GFAP (1:4000, rabbit, DAKO Z0334, RRID:AB_10013382), GFP (1:1000, chicken, Millipore AB16901, RRID:AB_90890), HA (1:1000, rat, Roche 11867423001, RRID:AB_390918), RFP (1:1000, rabbit, Rockland 600-401-379, RRID:AB_2209751). The next day, coverslips were washed 3 times with PBS, and Alexa Fluor conjugated secondaries (Invitrogen: Rb 594, RRID:AB_2534079; Ck 488 RRID:AB_2534096; Rat 594 RRID:AB_10561522) were diluted 1:1000 in blocking buffer containing 10% NGS and incubated for 2 hours at room temperature. After the incubation, the coverslips were washed 3 times and mounted onto glass slides (VWR) with Vectashield mounting media containing DAPI (Vector Laboratories).

Imaging was described as before(Stogsdill et al., 2017) and images were acquired at 21°C on either an Axioimager M1 (Zeiss) fluorescent microscope with an EC Plan-Neofluar 40× oil NA 1.3 objective (Immersol 518F) and an AxioCAM MRm camera, supported by the AxioVision SE64 software, and a Keyence BZ-X800 microscope with a Nikon S Plan Fluor ELWD ADM 40XC/0.60NA dry/oil objective and a 2/3-inch, 2.83 million pixel Peltier-cooled monochrome CCD (colorized with LC filter) camera, supporter by the BZ-X800 analyzer software. Images were acquired by a blinded investigator and analyzed with Sholl analysis in FIJI.

### CRISPR genome editing efficiency

To quantify the percent of frameshift inducing mutations, we cloned all 3 NL2 gRNAs and their respective PAM sequences upstream of and in frame with a GFP coding sequence. In HEK293T (ATCC, RRID:CVCL_4V93) cells we co-transfected the pX601-CMV-SaCas9-U6 gRNA plasmids (containing each NL2 gRNA) with the GFP target plasmid. Three days after transfection, the cells were harvested in 1X MPER (Thermo Scientific), incubated at 4C with end-over-end rotation for 10 minutes and then spun at 15,000 x g for 5 minutes. The supernatant was kept for downstream western blot analysis. GFP expression on the resulting western blot was quantified using ImageJ and normalized to GAPDH expression. In this assay, a reduction of GFP expression is interpreted to be genomic cutting by the corresponding gRNA.

### In vitro Sholl Analysis

Sholl Analysis was performed in FIJI (NIH, v1.53c) using the Sholl Analysis plug-in (v4.0.1). The number of intersections is measured with concentric circles at every 1µm starting from a radius of 10µm from the center of the nucleus. Resulting values were then analyzed in R (v4.0.0 and above) using an in-home pipeline which can be found here: https://github.com/Eroglu-Lab/In-Vitro-Sholl. The final graphs are the average of each biological replicate, not of each image. The line graphs were generated using the LOESS method (ggplot2 [Hadley Wickham], stat_smooth function) and the variance (standard error) is represented with the colored lines. The data were analyzed with a linear mixed effects model ANOVA, which accounts for the highly repetitious nature of Sholl analysis data by nesting each Radius measurement within the cell measured (Wilson et al., 2017).

### Postnatal astrocyte labeling by electroporation (PALE)

PALE was performed as previously described(Baldwin et al., 2021). Briefly, P0-P1 pups were placed on a glove on ice for < 5 minutes until unresponsive. 1-2µg (in 1-2µl total volume) of plasmid was loaded into a pulled glass pipette containing 1% fast green dye (Sigma F7252). The glass pipette was then inserted into one lateral ventricle of the mouse brain and DNA was injected. Electrodes were then positioned above the visual cortex (positive end) and below the jaw (negative end) and five 50ms pulses of 100V with an interpulse interval of 950ms were applied. Pups recovered on a heating pad and were then returned to the home cage with the dam. Each litter received a random assignment of experimental groups. All animals that recovered after surgery with no visible health issues were kept for the analysis. Animals were transcardially perfused with 4% PFA at P21.

### Immunohistochemistry

Mice were perfused transcardially with 4% PFA and were transferred to 4% PFA overnight before incubation with 30% sucrose in TBS. Coronal sections were collected on a cryostat (Leica) at either 40µm (iBioID, synaptic staining) or 100µm (territory volume and 3D Sholl). Sections were washed 3 times for 10 minutes on a shaker in 0.2%(40µm sections) or 0.4%(100µm sections) Triton-X in TBS (TBST). Sections were blocked in 10% Goat serum in TBST for 1 hour on the shaker at room temperature. Primary antibodies: PSD95 (1:300, rabbit, Invitrogen 51-6900, RRID:AB_2533914), Gephyrin (1:500, mouse, Synaptic Systems 147011, RRID:AB_2810215), RFP (1:1000, chicken, Rockland 600-901-379, RRID:AB_10704808, used for synaptic staining), Bassoon (1:500, guinea pig, Synaptic Systems 141318, RRID:AB_2927388), HA (1:200, rat, Roche 11867423001, RRID:AB_390918), RFP [for detecting mCherry CAAX in shRNA PALE experiments] (1:1000, rabbit, Rockland 600-401-379, RRID:AB_2209751), RFP nanobody 568 coupled [for detecting tdTomato in Ai14 mice for NL2 rescue PALE experiments] (1:100, Synaptic Systems,N0404-AF568-L, RRID:AB_2209751), Sox9 (1:1000, Rabbit, Millipore AB5535, RRID:AB_2239761), NeuN (1:1000, Mouse, Millipore MAB377, RRID:AB_2298772) were diluted in fresh blocking buffer and incubated either overnight (40µm) or for 3 nights (100µm). Alexa Fluor or Streptavidin conjugated fluorescent secondaries (Invitrogen: Rabbit 405+ RRID:AB_2890548; Mouse IgG1 488 RRID:AB_2535764, Chicken 568 RRID:AB_2534098, Guinea Pig 647 RRID:AB_2535867, Rat 488 RRID:AB_2534074, Rabbit 594 RRID:AB_2534095, Streptavidin Alexa Fluor conjugate (Thermo Fisher, S32357)) were diluted in blocking buffer at 1:200 and incubated on tissue for 3 hours at room temperature. DAPI was added in the final wash before mounting. Sections were mounted on glass slides with mounting media (90% Glycerol, 20 mM Tris pH 8.0, 0.5% n-Propyl gallate). Unless specified otherwise, images were acquired at 21°C on a Leica Stellaris 8 Confocal with a 63×/1.40 HC PL APO CS2 (11506350) oil objective (Leica F Immersion liquid [Oil] 11513859) and a Leica K5 sCMOS (14402610) camera, supported by the Leica Application Suite X (LASX, build 4.5.0.25531, RRID:SCR_013673, https://www.leica-microsystems.com/products/microscope-software/p/leica-las-x-ls/) software.

### Synapse quantification

Images for synapse quantification were acquired at 21°C on an Olympus FV3000 microscope with an Olympus Plan APO 60× 1.4 NA oil objective (Immoil-f30cc) and a 1.64X zoom, supported by the Olympus FV315-SW software. Images were captured in 1µm stacks with a 0.34µm step size. We used SynBot(Savage et al., 2024) (https://github.com/Eroglu-Lab/Syn_Bot) in FIJI to quantify the colocalization of Bassoon (visualized by Alexa 594) and Psd95 (excitatory, visualized by Alexa 405+) or Bassoon and Gepyhrin (inhibitory, visualized by Alexa 488) synapses for the entire image using manual thresholding. Using a custom macro in FIJI (https://github.com/Eroglu-Lab/Sakers-et-al-2025), we then cropped the binarized synaptic output images to draw an ROI around the labeled astrocyte and quantify the number of synaptic puncta within and outside the ROI. These synaptic puncta were normalized to the corresponding ROI to obtain a synapse density. The data were analyzed with a linear mixed effects ANOVA in R.

### Imaris territory volume and 3D Sholl analysis

100µm sections were imaged at 21°C on an Olympus FV3000 microscope with an Olympus Plan APO 60× 1.4 NA oil objective (Immoil-f30cc) and a 2× zoom, supported by the Olympus FV315-SW software. Single astrocytes (visualized by Alexa fluor 594 secondary antibodies) were captured in a frame in a Z stack at 0.5µm step size to capture the whole astrocyte. Imaris v9.9 (Bitplane) was used to reconstruct images using the filament tracer in 3D. We used two custom extensions to quantify filament complexity (3D Sholl analysis) every 5µm from the cell nucleus [note that this is different from 2D Sholl analysis where a measurement is taken every 1µm from the cell nucleus]. A linear mixed model followed by Dunnett’s post-hoc test was used to analyze the 3D Sholl data using a custom R script found at: https://github.com/Eroglu-Lab/In-vivo-Sholl-Analysis. The whole astrocyte territory volume was calculated using the Convex Hull function in Imaris on the cell surface. Analysis of territory volume was analyzed in R using a one-way repeated measures ANOVA to account for multiple images taken per animal, followed by Dunnett’s post-hoc test.

### Adeno-associated virus production

Purified AAV was produced as described previously(Baldwin et al., 2021). Briefly, HEK293T cells were transfected with pAd-DELTA F6, serotype plasmid AAV PHP.eB, and AAV plasmids (pZac2.1 Turbo ID constructs). Three days after transfection, cells were collected in 15 mM NaCl, 5 mM Tris-HCl, pH 8.5, and lysed with repeat freeze-thaw cycles followed by treatment with Benzonase (Novagen 70664) at 37°C for 30 minutes. Lysed cells were pelleted by centrifugation and the supernatant, containing AAV was applied to an Optiprep density gradient (Sigma D1556, 15%, 25%, 40%, and 60%) and centrifuged at 67,000 rpm using a Beckman Ti-70 rotor for 1 hour. The AAV-enriched fraction was isolated from between 40% and 60% iodixanol solution and concentrated by repeated washes with sterile PBS in an Amicon Ultra-15 filtration unit (NMWL: 100 kDa, Millipore UFC910008) to a final volume of ∼100 μL and aliquoted for storage at −80°C.

### In vivo BioID

Pregnant CD1 dams were ordered from Charles River Laboratories. On P1, pups were anesthetized by hypothermia and subsequently injected intracranially at a depth of 2mm with 1µl equal viral titers of either NL2-Turbo-BirA or Turbo-BirA as previously described. Mice were treated with 0.75mL of 5mM Biotin in sterile PBS via subcutaneous injections once a day from P18 to P20. On P21, mice were sacrificed with 200 mg/kg tribromoethanol (avertin) and transcardially perfused with TBS/Heparin for 5 minutes to wash out excess biotin. Subsequently, cortices were rapidly dissected in DPBS and flash-frozen in liquid nitrogen.

All solutions were made fresh using proteomics-grade chemicals and solvents. Brains were homogenized with approximately 12 strokes in a glass homogenizer and Teflon pestle in 1mL Lysis buffer [50mM Tris/HCl pH7.5, 150mM NaCl, 1mM EDTA with protease inhibitors (Roche)] on ice. An equal volume of 2X RIPA [50mM Tris/HCl pH7.5, 150mM NaCl, 1mM EDTA, 0.4% SDS, 2%TritonX100, 2% deoxycholate] was added to the lysates. Lysates were then sonicated (level 9) for 30 seconds (3 x 10s) total, followed by centrifugation at 15,000 RPM for 10 minutes at 4C. The supernatant was transferred to an ultracentrifuge 1.5mL tube (Beckman Coulter) and spun in a TLA-55 rotor (Beckman coulter) in an ultracentrifuge at 40,000 RPM for 30 minutes at 4C. Supernatants were transferred to a low-protein binding tube (Eppendorf) and SDS was added to a final concentration of 1%. Samples were then briefly vortexed and heated at 45C for 45 mins, cooled on ice, and spun at 15,000RPM for 30 minutes at 4C. 70 µl Neutravidin beads (ThermoFisher 29202) were prepared per condition by washing with 1mL of 2X RIPA two times. The samples were transferred to 15mL tubes (1 brain per IP, 3 brains per condition), and washed Neutravidin beads were added to the sample supernatant. 30ul of the supernatant was saved for western blot analysis. The beads and samples were incubated overnight at 4C with rotation. The next day beads were sequentially washed with 1mL of the following solutions for 10 minutes with rotation, followed by two-minute spins: 2% SDS x2, Wash buffer (1% TritonX100, 1% deoxycholate, 25 mM LiCl) x2, 1M NaCl x2, 50mM Ammonium bicarbonate x5. Elution buffer was prepared during the last washes by adding 1.2mg of biotin to 2X sample buffer [4% SDS, 20% glycerol, 0.1% beta-mercaptoethanol, 125mM Tris pH 6.8 to 20mL with keratin-free water]. 70ul of elution buffer was added to the beads and samples were then heated at 60C for 15 minutes to elute bound proteins. The sample was stored at –80C before mass spec analysis, and 5ul of the eluate was saved for western blotting. All washes and elution were done in a tissue culture hood, in sterile conditions to avoid keratin contamination.

### Mass spectrometry

Sample Preparation: The Duke Proteomics Core Facility (DPCF) received 6 samples (3 replicates each of two conditions). Samples were reduced with 10 mM dithiothreitol for 30 min at 80C and alkylated with 20 mM iodoacetamide for 30 min at room temperature. Next, they were supplemented with a final concentration of 1.2% phosphoric acid and 597 μL of S-Trap (Protifi) binding buffer (90% MeOH/100mM TEAB). Proteins were trapped on the S-Trap, digested using 20 ng/μl sequencing grade trypsin (Promega) for 1 hr at 47C, and eluted using 50 mM TEAB, followed by 0.2% FA, and lastly using 50% ACN/0.2% FA. All samples were then lyophilized to dryness and resuspended in 12μL 1%TFA/2% acetonitrile containing 12.5 fmol/μL yeast alcohol dehydrogenase (ADH_YEAST). A QC Pool was created from 3uL of each sample. All QC Pools were run periodically throughout the acquisition period.

Quantitative Analysis, Methods: Quantitative LC/MS/MS was performed on 3 μL of each sample, using a nanoAcquity UPLC system (Waters Corp) coupled to a Thermo Orbitrap Fusion Lumos high-resolution accurate mass tandem mass spectrometer (Thermo) via a nanoelectrospray ionization source. Briefly, the sample was first trapped on a Symmetry C18 20 mm × 180 μm trapping column (5 μl/min at 99.9/0.1 v/v water/acetonitrile), after which the analytical separation was performed using a 1.8 μm Acquity HSS T3 C18 75 μm × 250 mm column (Waters Corp.) with a 90-min linear gradient of 5 to 30% acetonitrile with 0.1% formic acid at a flow rate of 400 nanoliters/minute (nL/min) with a column temperature of 55C. Data collection on the Fusion Lumos mass spectrometer was performed in a data-dependent acquisition (DDA) mode of acquisition with a r=120,000 (@ m/z 200) full MS scan from m/z 375 – 1500 with a target AGC value of 2e5 ions. MS/MS scans were acquired at a Rapid scan rate (Ion Trap) with an AGC target of 5e3 ions and a max injection time of 100 ms. The total cycle time for MS and MS/MS scans was 2 sec. A 20s dynamic exclusion was employed to increase the depth of coverage. The total analysis cycle time for each sample injection was approximately 2 hours. Following 9 total UPLC-MS/MS analyses (excluding conditioning runs, but including 3 replicate QC Pool), data was imported into Proteome Discoverer 2.3 (Thermo Scientific Inc.), and analyses were aligned based on the accurate mass and retention time of detected ions (“features”) using Minora Feature Detector algorithm in Proteome Discoverer. Relative peptide abundance was calculated based on the area-under-the-curve (AUC) of the selected ion chromatograms of the aligned features across all runs. The MS/MS data was searched against the SwissProt M. musculus database (downloaded in Apr2017) and an equal number of reversed-sequence “decoys” for false discovery rate determination. Mascot Distiller and Mascot Server (v 2.5, Matrix Sciences) were utilized to produce fragment ion spectra and to perform the database searches. Database search parameters included fixed modifications on Cys (carbamidomethyl) and variable modifications on Meth (oxidation) and Asn and Gln (deamidation). Peptide Validator and Protein FDR Validator nodes in Proteome Discoverer were used to annotate the data at a maximum 1% protein false discovery rate.

#### Statistical analysis of iBioID data

In vivo BioID has an inherently high background in mass spectrometry due to the time necessary for biotin to accumulate in the system, and the diffuse nature of AMP-biotin. We controlled for this background using a soluble (non-bait) TurboID enzyme in both astrocytes and neurons, as this would accurately replicate the high background in bait conditions. Further, to account for differences in expression due to the size differenced of soluble TurboID and NL-tagged TurboID, we employed a robust-mean normalization which excludes the top and bottom 10% of signal intensities and mitigates the influences of outliers. The BioID datasets were generated from distinct viral constructs and cellular contexts, thus necessitating an enrichment-over-control design. To do so, we calculated the fold changes and used a two-tailed heteroscedastic t-test on log2-transformed data comparing the NL2 and Turbo BioID groups, and performed FDR correction to obtain high-confidence putative interactors.

### Western blotting

iBioID: 5ul of IP eluate and 30ul of sonicated lysate were run on a 4-15% Mini-Protean precast gel (BioRad). After transfer, membranes were blocked in Odyssey blocking buffer (LI-COR) diluted in PBS (1:1) followed by incubation in the same buffer with Alexa 680 conjugated streptavidin (Invitrogen). Membranes were imaged on a LI-COR Odyssey CLx. Densitometry analysis was performed in ImageJ (NIH v1.54f)

All other blots: Equal protein (measured by Bicinchoninic Acid Protein Assay (BCA, Pierce)) was loaded on a 4-15% Mini-Protean precast gel (BioRad). In the case of immunoprecipitations, input samples were loaded as an equal percentage of total sample homogenate, and the entirety of the IP eluate was run per lane. Gels were transferred using a Power Blotter System (Invitrogen) and then blocked in a Blocking Buffer for Fluorescent Western Blotting (Rockland, MB-070) diluting in PBS (1:1). The following primary antibodies were used for blotting: Rabbit anti-Nedd4l (Cell Signaling Technology, 4013S, 1:1000, RRID:AB_1904063), Rabbit anti-V5 (Cell Signaling Technology, 13202S, 1:500, RRID:AB_2687461), Chicken anti-HA (Aves Labs, ET-HA100, 1:2000, RRID:AB_2313511), Rabbit anti-GFP (Novus Biologicals, NB600-308, 1:1000, RRID:AB_10003058), Mouse anti-Gapdh (Abcam, ab8245, 1:2500, RRID:AB_2107448), Mouse anti-Actin (Sigma-Aldrich, A5441, 1:2000, RRID:AB_476744) diluted in blocking buffer. LI-COR 680 or 800 fluorescent secondary antibodies (Chicken 800 RRID:AB_10974977; Rabbit 800 RRID:AB_621848; Mouse 800 RRID:AB_621842, Rabbit 680 RRID:AB_621845, Chicken 680 RRID:AB_1850018) were diluted 1:2500 in blocking buffer and blots were imaged on a LI-COR Odyssey CLx.

### Lentivirus production and transduction

8-10 million 293T cells were plated in a T-75 culture flask and transfected using Xtremegene HP Transfection Reagent (Roche, 06366236001) with the following plasmids: pLKO.1 puromycin-resistant shRNA plasmids (described above), VSVG (an envelope plasmid), and dR8.91 packaging plasmid. One day post-transfection, the media was replaced with astrocyte growth media, and lentivirus was collected on days 2 and 3 and filtered through a 40µm syringe filter. Media was tested for lentiviral particles using Lenti-X Go Stix (Takara, 631280). Astrocytes were treated with 500µl of lentiviral media supernatant, and 1µg/mL polybrene on DIV8. On DIV10, astrocytes were treated with 1µg/mL puromycin every other day until a non-transduced well was completely dead. Cells were lysed for qPCR as described below. Cells collected for western blot were collected in T-PER (Thermo Scientific, 78510) and protein was measured using a BCA protein assay (Thermo Scientific, 23227).

### Gene ontology analysis

Gene ontology analysis was performed in R using the package clusterProfiler v4.6.2(Wu et al., 2021). Proteins from iBioID lists that were at least 1.5 fold enriched at a p-value of <0.05, compared to respective controls described in legends, were subjected to Biological Processes enrichment. Resulting output was reduced using the simplify function in clusterProfiler. The top 5-10 categories (sorted by adjusted p-value) were plotted in R. The entire enrichment can be found in Supplemental Dataset 2.

### Immunoprecipitation

4 million 293T cells (ATCC, RRID:CVCL_4V93) (passage #8-15) were plated in 10cm dishes and then transfected the next day with 2.5µg of each plasmid using the Xtremegene HP Transfection Reagent (Roche, 06366236001) according to the manufacturer’s protocol. 48 hours post transfection cells were washed with 1X TBS supplemented with calcium and magnesium and pelleted. Cell pellets were resuspended in 400µl of lysis buffer (25mM Tris pH 7.4, 1mM Calcium Chloride, 1mM Magnesium Chloride, 150mM Sodium Chloride, 0.5% NP-40, 1X EDTA-free protease inhibitor cocktail (Roche)) and centrifuged at 20,000xg for 10 minutes. 5% of the supernatant was saved for input samples, and the remaining lysate was diluted with lysis buffer without NP-40 for IP. Lysate was added to 30µl of magnetic anti-HA beads (Pierce, 88836) overnight at 4C with rotation. Beads were collected on a magnetic rack and washed 3 times with wash buffer (25mM Tris pH 7.4, 1mM Calcium Chloride, 1mM Magnesium Chloride, 150mM Sodium Chloride, 0.1% NP-40, 1X EDTA-free protease inhibitor cocktail (Roche)). Beads were eluted with 2X Laemmli buffer (Bio-Rad, 1610737) at 42C for 15 minutes and then denatured with 5% β-mercaptoethanol at 45C for 45 minutes before loading on SDS-PAGE. Bands were quantified via densitometry (ImageJ). Co-IP bands were normalized to the IP band of bait protein (Figure 6, HA-NLs). Paired Student’s t-tests were used to compare differences across condition, using biological replicates as pairs.

### Ubiquitination assay

293T cells were transfected as indicated in the immunoprecipitation subsection. 48 hours post-transfection, cells were treated with 25µM MG-132 (Sigma-Aldrich, 474790) for 2-4 hours to block the proteasome. Cells were washed 3 times in ice-cold 1X TBS supplemented with magnesium, calcium, and N-ethylmaleimide (NEM) (Sigma-Aldrich, E3876) to block deubiquitinases and then pelleted. Cells were resuspended in 1X RIPA buffer with SDS (50 mM Tris-HCl pH 7.4, 150 mM NaCl, 1% NP-40, 0.5% Sodium Deoxycholate, 1% Sodium dodecyl sulfate, supplemented with 1X protease inhibitor and 20mM NEM) and then boiled for 5 minutes to disrupt protein-protein interactions. The lysates were diluted 10X with 1X RIPA without SDS and incubated on a rotator at 4C for 1 hour. The lysates were then pelleted at 15,000xg for 15 minutes and the supernatant was added to magnetic anti-HA beads, and 5% of the supernatant was collected for input fraction. Elution was performed identically to the immunoprecipitation subsection above. For ubiquitination assay in primary rat astrocytes, 5 million DIV7 astrocytes (prepared as described above) were nucleofected with the Basic Glial Cells Nucleofection Kit (Lonza) and 5ug of each plasmid. 24 hours after nucleofection, the media was changed to remove dead cells and nucleofection solution. Ubiquitination IP was performed 3 days after nucleofection in the same manner as 293T cells.

### Degradation assay

350,000 HEK 293T cells were plated into wells of a 6-well plate and then transfected one day later with 1µg of each plasmid via Xtremegene HP Transfection Reagent reagent (Roche #06366236001). 48 hours later cells were treated with cycloheximide (100µg/mL). At each time point, the collected cells were washed with 1X TBS with magnesium and calcium, pelleted, and then resuspended in 1X RIPA (Sigma-Aldrich) supplemented with protease inhibitors. Lysates were incubated at 4C with rotation for 40 minutes, centrifuged at 20,000xg at 4C and supernatants were saved for protein quantitation via BCA. 5µg of total protein was loaded on a 4-15% SDS-PAGE. Three independent cultures were used for statistical analysis.

### Statistics

Experiments were randomized where applicable and analyses were run by an experimenter blind to the condition. All in vitro experiments were performed at least 3 times (3 different cultures), with 20+ cells imaged per replicate. Note that some conditions were plotted across several figures (i.e. shNL2 in Figure S1, Figure 1; shNL2 + HA-NL2 Figure 1D and F and S1, and shNL2 in Figure 5), as experiments were done in parallel. All analyses were done in R (v4.0.0 and above). An R script with reproducible data can be found at the following links for both in vitro analysis (https://github.com/Eroglu-Lab/In-Vitro-Sholl) and in vivo analysis (https://github.com/Eroglu-Lab/In-vivo-Sholl-Analysis). Sholl analysis statistical analysis was performed using a linear mixed effects ANOVA which accurately accounts for the multilevel and repetitive nature of the data(Wilson et al., 2017). Statistical analyses were performed using N of independent culture or animal rather than image to avoid pseudoreplication artifacts. Specific details on sample size, and test applied can be found in the figure legends and Methods subsections and in Table S1.

## Data availability statement

The mass spectrometry proteomics data have been deposited to the ProteomeXchange Consortium via the PRIDE(Perez-Riverol et al., 2022) partner repository with the dataset identifier PXD049185. All data in this manuscript can be found in the attached Source Data file, and all original images can be accessed via Duke Digital Repository: https://doi.org/10.7924/r4r500.

## Conflict of Interest Statement

The authors declare no competing financial interests.

## Author contributions

Conceptualization: Kristina Sakers and Cagla Eroglu; Investigation: Kristina Sakers, Juan J. Ramirez, Nimrod Elazar, Leykashree Nagendren; Methodology: Kristina Sakers, Cagla Eroglu; Data curation: Kristina Sakers; Formal analysis: Kristina Sakers, Erik Soderblom; Writing: Kristina Sakers, Cagla Eroglu. Project administration: Kristina Sakers; Supervision: Cagla Eroglu; Funding acquisition: Cagla Eroglu, Kristina Sakers, Nimrod Elazar and Juan J. Ramirez.

## FIGURE LEGENDS

**Supplemental Figure 1:**
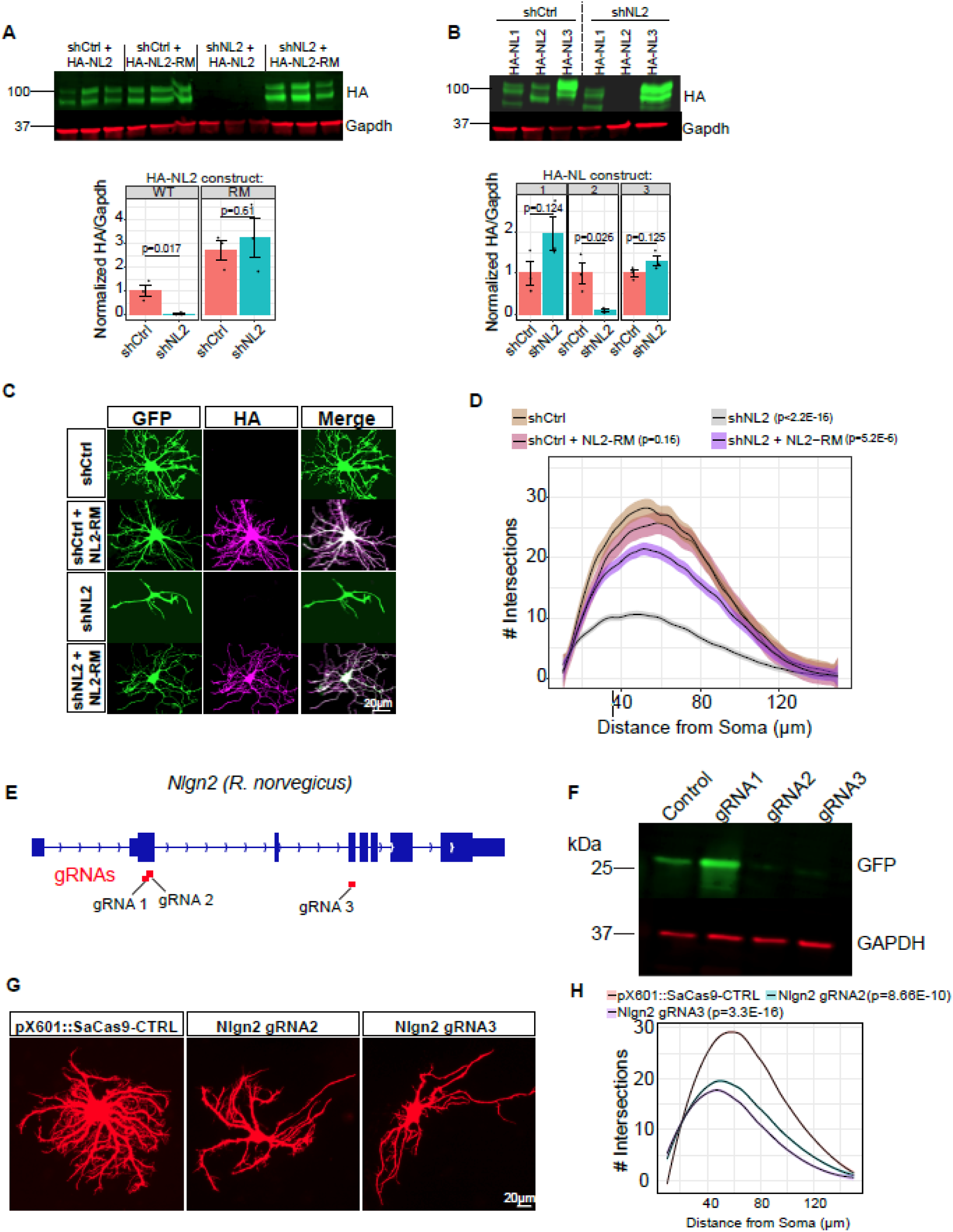
A) Western blots of HA-NL2 expressed in HEK293T cells. HEK293T cells were transfected with shCtrl or shNL2, and either HA-NL2 or HA-NL2-RM (Rescue mutant) to assess the efficacy of shNL2. HA and Gapdh bands were quantified via densitometry. HA was normalized to Gapdh to produce the graph below. HA-NL2 is reduced by >90% with shNL2. shNL2 has no effect on the HA-NL2-RM construct. Statistics are the result of a Student’s t-test. B) Western blots of HA-NL2 expressed in HEK293T cells. HEK293T cells were transfected with shCtrl or shNL2 and either HA-NL1, HA-NL2 or HA-NL3 to assess the specificity of shNL2. HA and Gapdh bands were quantified via densitometry. HA was normalized to Gapdh to produce the graph below. HA-NL2 is specifically targeted by shNL2. shNL2 results in non-significant increases in HA-NL1 and HA-NL3. Statistics are the result of a Student’s t-test. C) Representative images of astrocytes cultured on neurons. Astrocytes were transfected with either shCtrl or shNL2, alone or with HA-NL2-RM (Rescue mutant). D) Sholl analysis of images represented in A. Linear-mixed model followed by ANOVA reveals the main effect of condition F(3, 286)=75.01, p<2.2E-16. P-values represent post hoc Dunnett’s comparison to shCtrl. N = 53-96 cells per condition, across 3 biological replicates (independent culture experiments). Shaded area indicates standard error. E) *Nlgn2* sequence illustrating location of the gRNAs using in D and E. F) Western blot of GFP translation assay. The target sequence of the gRNAs is cloned upstream and in-frame with GFP. gRNAs are co-expressed and then lysates are collected and run in equal protein load on an SDS-PAGE. The western blot was probed for GFP and GAPDH, for normalization. G) Representative images of astrocytes cultured on neurons. Astrocytes were transfected with pX601 (U6 gRNA GfaABC1D Cas9 T2A mCherry) prior to culturing on neurons. H) Sholl analysis of images represented in C. Linear-mixed model followed by ANOVA reveals the main effect of condition F(2, 185) = 37.0175, p=3E-14. P-values represent post hoc Tukey’s comparison to shCtrl. N = 40 cells per condition, across 3 biological replicates (independent culture experiments). Shaded area indicates standard error.

**Supplemental Figure 2:**
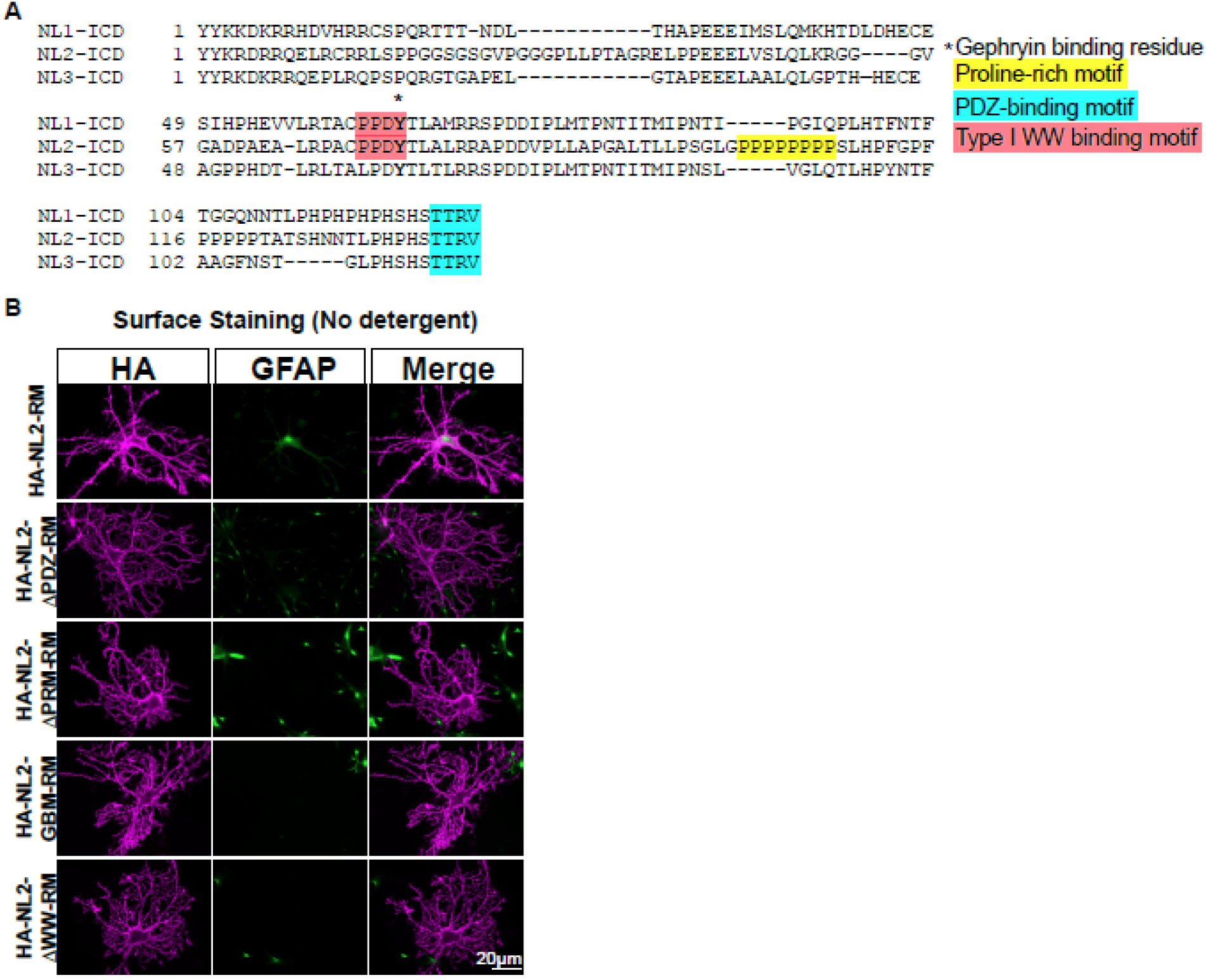
A) Mouse Neuroligin intracellular domain alignment, highlighting relevant protein sequence motifs. B) Representative images of surface-stained astrocyte/neuron co-cultures. Cells were fixed with cold 4% PFA for 5 minutes, and triton was omitted from blocking steps to avoid intracellular staining. GFAP was used as a negative control. HA (magenta) signal shows membrane localization as expected. GFAP signal is largely absent in the absence of detergent.

**Supplemental Figure 3:**
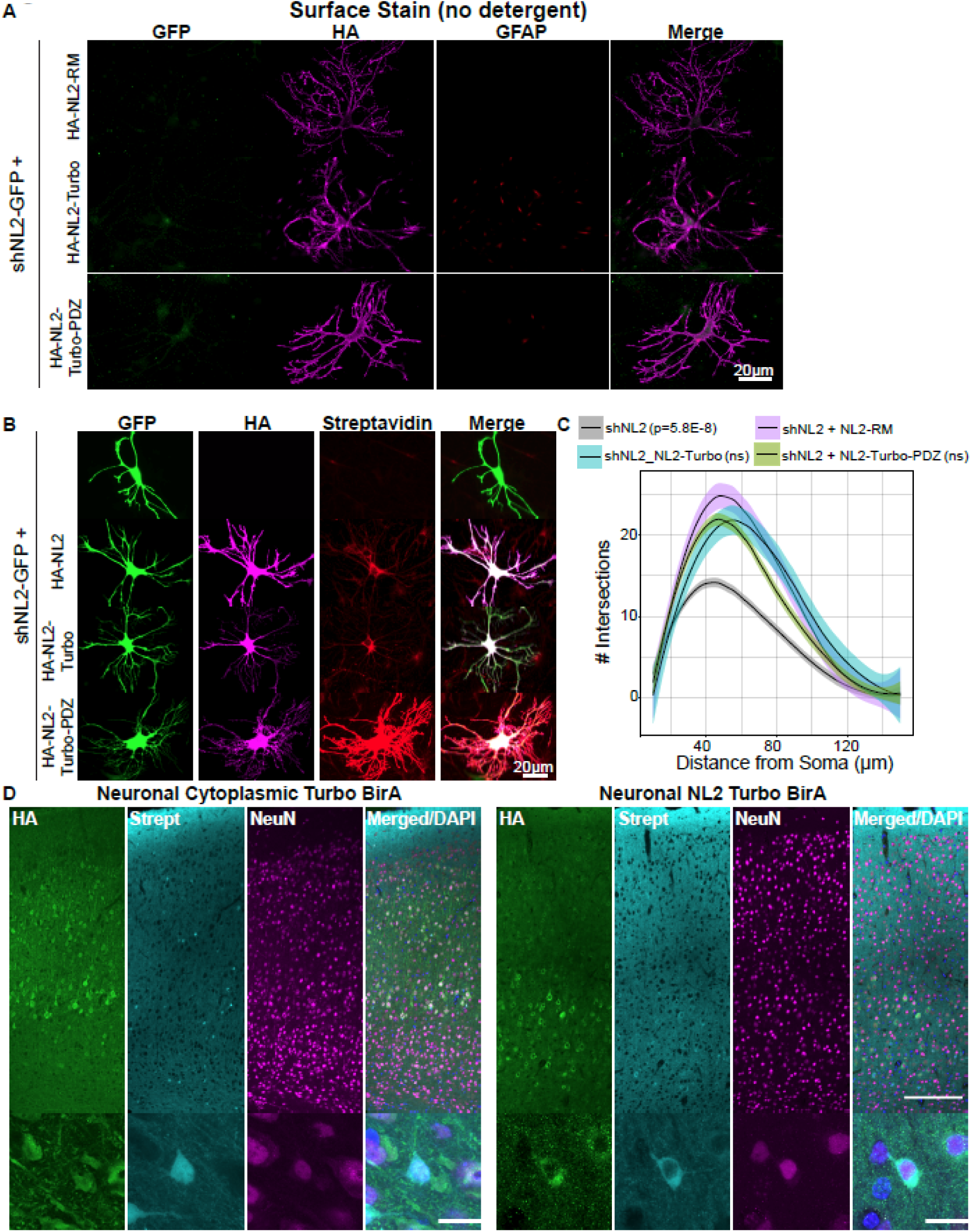
A) Representative images of surface-stained astrocyte/neuron co-cultures. Cells were fixed with cold 4% PFA for 5 minutes, and triton was omitted from blocking steps to avoid intracellular staining. GFP and GFAP were used as negative controls. HA (magenta) signal shows membrane localization as expected. B) Representative images demonstrating effective biotinylation (Streptavidin Alexa 594, red) in HA-NL2-Turbo-PDZ but not in HA-NL2 (negative control) or HA-NL2-Turbo. Experiments were performed at least twice with at least 20 cells per condition. C) Sholl analysis quantification of images in B. Linear-mixed model reveals the main effect of condition, F(1,3)=11.01, p=1.4E-6. P-values represent Dunnett’s post-hoc with respect to shNL2 + NL2-RM. NS = not significant in Dunnett’s. N = 14-56 cells across 3 biological replicates (independent culture experiments). Shaded area indicates standard error. D). Representative cortical images of Neuronal Turbo BirA injected animals. Stained with Streptavidin-594 (Strept), HA and NeuN to determine neuronal specificity. Scale bars: column image = 100µm; inset = 20µm.

**Supplemental Figure 4:**
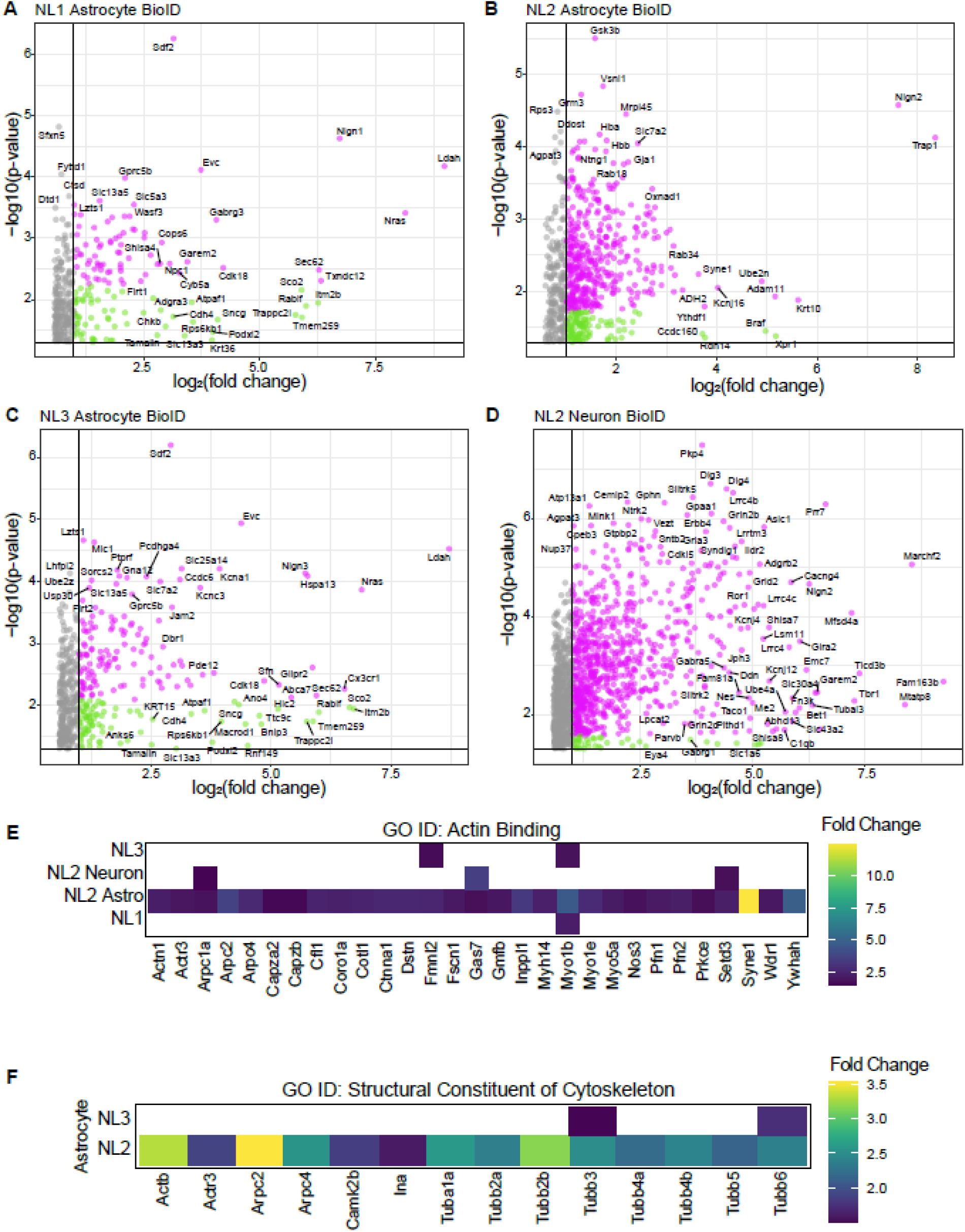
A-D) Volcano plots of all proteins identified in the iBioID experiments. Pink dots indicate proteins enriched over cytoplasmic control with a fold change > 1.5 and an FDR-corrected p-value < 0.05. Green dots indicate proteins enriched over cytoplasmic control with a fold change > 1.5 and a p-value < 0.05. E-F) Expression of genes related to Actin Binding (E) or Structural Constituent of Cytoskeleton (F) from Gene Ontology analysis of iBioID shows that astrocytic NL2 significantly interacts with cytoskeletal genes, compared to astrocyte NL1/NL3 or neuronal NL2.

**Supplemental Figure 5:**
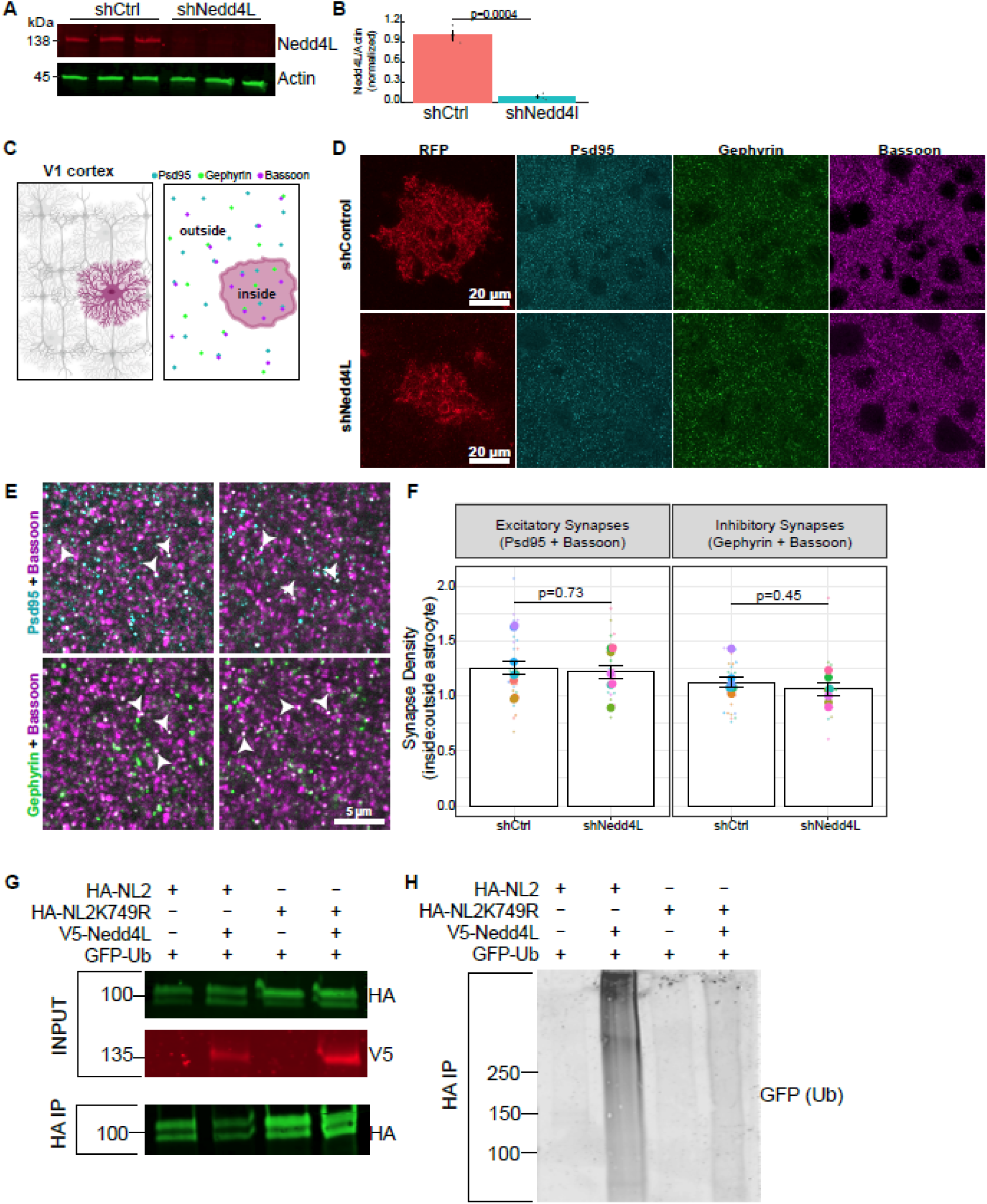
A) Western blots of astrocytes treated with lentivirus expressing shCtrl (scrambled shRNA) or shNedd4l. B) Densitometry quantification [ImageJ]. The p value represents the results of the Student’s t-test. C) Cartoon of synapse analysis. Sparsely labeled astrocytes (shControl mCherry CAAX; shNedd4l mCherry CAAX) were imaged in P21 mouse V1 cortex. Synapses (colocalized Psd95 + Bassoon [excitatory synapses] or Gephyrin + Bassoon [inhibitory synapses]) were quantified within the astrocyte territory (inside) and outside of the astrocyte territory (outside). D) Representative images of mCherry-labeled astrocytes (RFP antibody), with Psd95 (cyan), Gephyrin (green), Bassoon (magenta) staining. E) Magnified image showing colocalized synapse puncta for excitatory (Psd95 + Bassoon), and inhibitory (Gephyrin and Bassoon) synapses. Colocalized puncta are shown in white, with examples indicated with arrows. F) Quantification of synapse density within labeled astrocyte to the outside unlabeled astrocyte territory. Large dots represent individual animals, with small dots representing each image from an animal. P-values indicate main effect of linear mixed effects ANOVA. Excitatory synapses: F(1,13) = 0.12, p=0.73. Inhibitory synapses: F(1,13) = 0.61, p=0.45. 7-8 animals were used per condition, with 4-8 images taken per animal. G-H) Western blot from HEK cell lysates and HA-NL2 IPs (G) shows NL2 ubiquitination (H) with Nedd4l overexpression, which is substantially reduced when NL2 is mutated at K749. GFP-tagged ubiquitin was overexpressed in these cells to visualize ubiquitinated NL2 in the immunoprecipitation.

**Supplemental Dataset 1: In vivo BioID results.** All proteins significantly enriched in Astrocytic NL1 (sheet 1), Astrocytic NL2 (sheet 2), Astrocytic NL3 (sheet 3), and Neuronal NL (sheet 4) iBioID compared to Astrocytic TurboID alone.

**Supplemental Dataset 2: iBioID Gene Ontology output.** All biological process gene ontology results. Column 1 indicates the iBioID list to which the categories correspond to.

**Supplemental Table 1: Detailed statistical analyses for each figure panel.**

## Notes

### Competing Interest Statement

The authors have declared no competing interest.

### Summary of Updates

Added new data figures (S1A-B); revised text for clarity; corrected an error in sholl analysis statistics throughout the manuscript.

## REFERENCES

1. Ackerman, S.D., N.A. Perez-Catalan, M.R. Freeman, and C.Q. Doe. 2021. Astrocytes close a motor circuit critical period. Nature. 592:414–420. doi:10.1038/s41586-021-03441-2.

2. Altas, B., H.-J. Rhee, A. Ju, H.C. Solís, S. Karaca, J. Winchenbach, O. Kaplan-Arabaci, M. Schwark, M.C. Ambrozkiewicz, C. Lee, L. Spieth, G.L. Wieser, V.K. Chaugule, I. Majoul, M.A. Hassan, R. Goel, S.M. Wojcik, N. Koganezawa, K. Hanamura, D. Rotin, A. Pichler, M. Mitkovski, L. de Hoz, A. Poulopoulos, H. Urlaub, O. Jahn, G. Saher, N. Brose, J. Rhee, and H. Kawabe. 2024. Nedd4-2-dependent regulation of astrocytic Kir4.1 and Connexin43 controls neuronal network activity. J. Cell Biol. 223:e201902050. doi:10.1083/jcb.201902050.

3. Babaev, O., P. Botta, E. Meyer, C. Müller, H. Ehrenreich, N. Brose, A. Lüthi, and D. Krueger-Burg. 2016. Neuroligin 2 deletion alters inhibitory synapse function and anxiety-associated neuronal activation in the amygdala. Neuropharmacology. 100:56–65. doi:10.1016/j.neuropharm.2015.06.016.

4. Baldwin, K.T., K.K. Murai, and B.S. Khakh. 2023. Astrocyte morphology. Trends Cell Biol. S0962-8924(23)00204–0. doi:10.1016/j.tcb.2023.09.006.

5. Baldwin, K.T., C.X. Tan, S.T. Strader, C. Jiang, J.T. Savage, X. Elorza-Vidal, X. Contreras, T. Rülicke, S. Hippenmeyer, R. Estévez, R.-R. Ji, and C. Eroglu. 2021. HepaCAM controls astrocyte self-organization and coupling. Neuron. 109:2427–2442.e10. doi:10.1016/j.neuron.2021.05.025.

6. Barrow, S.L., J.R. Constable, E. Clark, F. El-Sabeawy, A.K. McAllister, and P. Washbourne. 2009. Neuroligin1: a cell adhesion molecule that recruits PSD-95 and NMDA receptors by distinct mechanisms during synaptogenesis. Neural Develop. 4:17. doi:10.1186/1749-8104-4-17.

7. Binda, C.S., Y. Nakamura, J.M. Henley, and K.A. Wilkinson. 2019. Sorting nexin 27 rescues neuroligin 2 from lysosomal degradation to control inhibitory synapse number. Biochem. J. 476:293–306. doi:10.1042/BCJ20180504.

8. Boisvert, M.M., G.A. Erikson, M.N. Shokhirev, and N.J. Allen. 2018. The Aging Astrocyte Transcriptome from Multiple Regions of the Mouse Brain. Cell Rep. 22:269–285. doi:10.1016/j.celrep.2017.12.039.

9. Booth, H.D.E., W.D. Hirst, and R. Wade-Martins. 2017. The Role of Astrocyte Dysfunction in Parkinson’s Disease Pathogenesis. Trends Neurosci. 40:358–370. doi:10.1016/j.tins.2017.04.001.

10. Branon, T.C., J.A. Bosch, A.D. Sanchez, N.D. Udeshi, T. Svinkina, S.A. Carr, J.L. Feldman, N. Perrimon, and A.Y. Ting. 2018. Efficient proximity labeling in living cells and organisms with TurboID. Nat. Biotechnol. 36:880–887. doi:10.1038/nbt.4201.

11. Bruce, M.C., V. Kanelis, F. Fouladkou, A. Debonneville, O. Staub, and D. Rotin. 2008. Regulation of Nedd4-2 self-ubiquitination and stability by a PY motif located within its HECT-domain. Biochem. J. 415:155–163. doi:10.1042/BJ20071708.

12. Bushong, E.A., M.E. Martone, and M.H. Ellisman. 2004. Maturation of astrocyte morphology and the establishment of astrocyte domains during postnatal hippocampal development. Int. J. Dev. Neurosci. 22:73–86. doi:10.1016/j.ijdevneu.2003.12.008.

13. Bushong, E.A., M.E. Martone, Y.Z. Jones, and M.H. Ellisman. 2002. Protoplasmic astrocytes in CA1 stratum radiatum occupy separate anatomical domains. J. Neurosci. Off. J. Soc. Neurosci. 22:183–192. doi:10.1523/JNEUROSCI.22-01-00183.2002.

14. Chanda, S., W.D. Hale, B. Zhang, M. Wernig, and T.C. Südhof. 2017. Unique versus Redundant Functions of Neuroligin Genes in Shaping Excitatory and Inhibitory Synapse Properties. J. Neurosci. Off. J. Soc. Neurosci. 37:6816–6836. doi:10.1523/JNEUROSCI.0125-17.2017.

15. Chen, H.I., A. Einbond, S.J. Kwak, H. Linn, E. Koepf, S. Peterson, J.W. Kelly, and M. Sudol. 1997. Characterization of the WW domain of human yes-associated protein and its polyproline-containing ligands. J. Biol. Chem. 272:17070–17077. doi:10.1074/jbc.272.27.17070.

16. Chih, B., H. Engelman, and P. Scheiffele. 2005. Control of excitatory and inhibitory synapse formation by neuroligins. Science. 307:1324–1328. doi:10.1126/science.1107470.

17. Chih, B., L. Gollan, and P. Scheiffele. 2006. Alternative splicing controls selective trans-synaptic interactions of the neuroligin-neurexin complex. Neuron. 51:171–178. doi:10.1016/j.neuron.2006.06.005.

18. Clarke, L.E., S.A. Liddelow, C. Chakraborty, A.E. Münch, M. Heiman, and B.A. Barres. 2018. Normal aging induces A1-like astrocyte reactivity. Proc. Natl. Acad. Sci. 115:E1896–E1905. doi:10.1073/pnas.1800165115.

19. Concordet, J.-P., and M. Haeussler. 2018. CRISPOR: intuitive guide selection for CRISPR/Cas9 genome editing experiments and screens. Nucleic Acids Res. 46:W242–W245. doi:10.1093/nar/gky354.

20. Endo, F., A. Kasai, J.S. Soto, X. Yu, Z. Qu, H. Hashimoto, V. Gradinaru, R. Kawaguchi, and B.S. Khakh. 2022. Molecular basis of astrocyte diversity and morphology across the CNS in health and disease. Science. 378:eadc9020. doi:10.1126/science.adc9020.

21. Farhy-Tselnicker, I., M.M. Boisvert, H. Liu, C. Dowling, G.A. Erikson, E. Blanco-Suarez, C. Farhy, M.N. Shokhirev, J.R. Ecker, and N.J. Allen. 2021. Activity-dependent modulation of synapse-regulating genes in astrocytes. eLife. 10:e70514. doi:10.7554/eLife.70514.

22. Farizatto, K.L.G., and K.T. Baldwin. 2023. Astrocyte-synapse interactions during brain development. Curr. Opin. Neurobiol. 80:102704. doi:10.1016/j.conb.2023.102704.

23. Giannone, G., M. Mondin, D. Grillo-Bosch, B. Tessier, E. Saint-Michel, K. Czöndör, M. Sainlos, D. Choquet, and O. Thoumine. 2013. Neurexin-1β Binding to Neuroligin-1 Triggers the Preferential Recruitment of PSD-95 versus Gephyrin through Tyrosine Phosphorylation of Neuroligin-1. Cell Rep. 3:1996–2007. doi:10.1016/j.celrep.2013.05.013.

24. Goel, K., and J.E. Ploski. 2022. RISC-y Business: Limitations of Short Hairpin RNA-Mediated Gene Silencing in the Brain and a Discussion of CRISPR/Cas-Based Alternatives. Front. Mol. Neurosci. 15. doi:10.3389/fnmol.2022.914430.

25. Golf, S.R., J.H. Trotter, J. Wang, G. Nakahara, X. Han, M. Wernig, and T.C. Südhof. 2025. Deletion of Neuroligins from Astrocytes Does Not Detectably Alter Synapse Numbers or Astrocyte Cytoarchitecture by Maturity. eLife. 12. doi:10.7554/eLife.87589.3.

26. Gupta, S., J.A. Stamatoyannopoulos, T.L. Bailey, and W.S. Noble. 2007. Quantifying similarity between motifs. Genome Biol. 8:R24. doi:10.1186/gb-2007-8-2-r24.

27. Gutmann, D.H., A. Loehr, Y. Zhang, J. Kim, M. Henkemeyer, and A. Cashen. 1999. Haploinsufficiency for the neurofibromatosis 1 (NF1) tumor suppressor results in increased astrocyte proliferation. Oncogene. 18:4450–4459. doi:10.1038/sj.onc.1202829.

28. Halff, E.F., B.R. Szulc, F. Lesept, and J.T. Kittler. 2019. SNX27-Mediated Recycling of Neuroligin-2 Regulates Inhibitory Signaling. Cell Rep. 29:2599–2607.e6. doi:10.1016/j.celrep.2019.10.096.

29. Hatstat, A.K., M.D. Pupi, and D.G. McCafferty. 2021. Predicting PY motif-mediated protein-protein interactions in the Nedd4 family of ubiquitin ligases. PLOS ONE. 16:e0258315. doi:10.1371/journal.pone.0258315.

30. Holt, L.M., R.D. Hernandez, N.L. Pacheco, B. Torres Ceja, M. Hossain, and M.L. Olsen. 2019. Astrocyte morphogenesis is dependent on BDNF signaling via astrocytic TrkB.T1. eLife. 8:e44667. doi:10.7554/eLife.44667.

31. Hunker, A.C., and L.S. Zweifel. 2020. Protocol to Design, Clone, and Validate sgRNAs for In Vivo Reverse Genetic Studies. STAR Protoc. 1:100070. doi:10.1016/j.xpro.2020.100070.

32. Ilsley, J.L., M. Sudol, and S.J. Winder. 2002. The WW domain: linking cell signalling to the membrane cytoskeleton. Cell. Signal. 14:183–189.

33. Irie, M., Y. Hata, M. Takeuchi, K. Ichtchenko, A. Toyoda, K. Hirao, Y. Takai, T.W. Rosahl, and T.C. Südhof. 1997. Binding of Neuroligins to PSD-95. Science. 277:1511–1515. doi:10.1126/science.277.5331.1511.

34. Jamain, S., H. Quach, C. Betancur, M. Råstam, C. Colineaux, I.C. Gillberg, H. Söderström, B. Giros, M. Leboyer, C. Gillberg, and T. Bourgeron. 2003. Mutations of the X-linked genes encoding neuroligins NLGN3 and NLGN4 are associated with autism. Nat. Genet. 34:27–29. doi:10.1038/ng1136.

35. Kang, Y., Y. Ge, R.M. Cassidy, V. Lam, L. Luo, K.-M. Moon, R. Lewis, R.S. Molday, R.O.L. Wong, L.J. Foster, and A.M. Craig. 2014. A Combined Transgenic Proteomic Analysis and Regulated Trafficking of Neuroligin-2. J. Biol. Chem. 289:29350–29364. doi:10.1074/jbc.M114.549279.

36. Kasanov, J., G. Pirozzi, A.J. Uveges, and B.K. Kay. 2001. Characterizing Class I WW domains defines key specificity determinants and generates mutant domains with novel specificities. Chem. Biol. 8:231–241.

37. Kato, Y., M. Ito, K. Kawai, K. Nagata, and M. Tanokura. 2002. Determinants of Ligand Specificity in Groups I and IV WW Domains as Studied by Surface Plasmon Resonance and Model Building *. J. Biol. Chem. 277:10173–10177. doi:10.1074/jbc.M110490200.

38. Kato, Y., K. Nagata, M. Takahashi, L. Lian, J.J. Herrero, M. Sudol, and M. Tanokura. 2004. Common mechanism of ligand recognition by group II/III WW domains: redefining their functional classification. J. Biol. Chem. 279:31833–31841. doi:10.1074/jbc.M404719200.

39. Kirkeby, S., D. Moe, T.C. Bøg-Hansen, and C.J.F. van Noorden. 1993. Biotin carboxylases in mitochondria and the cytosol from skeletal and cardiac muscle as detected by avidin binding. Histochemistry. 100:415–421. doi:10.1007/BF00267821.

40. Lavialle, M., G. Aumann, E. Anlauf, F. Pröls, M. Arpin, and A. Derouiche. 2011. Structural plasticity of perisynaptic astrocyte processes involves ezrin and metabotropic glutamate receptors. Proc. Natl. Acad. Sci. U. S. A. 108:12915–12919. doi:10.1073/pnas.1100957108.

41. Lee, D.-E., J.E. Yoo, J. Kim, S. Kim, S. Kim, H. Lee, and H. Cheong. 2020. NEDD4L downregulates autophagy and cell growth by modulating ULK1 and a glutamine transporter. Cell Death Dis. 11:1–17. doi:10.1038/s41419-020-2242-5.

42. Lee, K.Y., K.A. Jewett, H.J. Chung, and N.-P. Tsai. 2018. Loss of fragile X protein FMRP impairs homeostatic synaptic downscaling through tumor suppressor p53 and ubiquitin E3 ligase Nedd4-2. Hum. Mol. Genet. 27:2805–2816. doi:10.1093/hmg/ddy189.

43. Lee, Y., A. Messing, M. Su, and M. Brenner. 2008. GFAP promoter elements required for region-specific and astrocyte-specific expression. Glia. 56:481–493. doi:10.1002/glia.20622.

44. Letellier, M., Z. Szíber, I. Chamma, C. Saphy, I. Papasideri, B. Tessier, M. Sainlos, K. Czöndör, and O. Thoumine. 2018. A unique intracellular tyrosine in neuroligin-1 regulates AMPA receptor recruitment during synapse differentiation and potentiation. Nat. Commun. 9:3979. doi:10.1038/s41467-018-06220-2.

45. Li, J., T.G. Miramontes, T. Czopka, and K.R. Monk. 2024. Synaptic input and Ca2+ activity in zebrafish oligodendrocyte precursor cells contribute to myelin sheath formation. Nat. Neurosci. doi:10.1038/s41593-023-01553-8.

46. Liu, X., H. Zhang, B. Zhang, J. Tu, X. Li, and Y. Zhao. 2021. Nedd4-2 haploinsufficiency in mice causes increased seizure susceptibility and impaired Kir4.1 ubiquitination. Biochim. Biophys. Acta Mol. Basis Dis. 1867:166128. doi:10.1016/j.bbadis.2021.166128.

47. Loh, K.H., P.S. Stawski, A.S. Draycott, N.D. Udeshi, E.K. Lehrman, D.K. Wilton, T. Svinkina, T.J. Deerinck, M.H. Ellisman, B. Stevens, S.A. Carr, and A.Y. Ting. 2016. Proteomic Analysis of Unbounded Cellular Compartments: Synaptic Clefts. Cell. 166:1295–1307.e21. doi:10.1016/j.cell.2016.07.041.

48. Minegishi, S., T. Ishigami, T. Kino, L. Chen, R. Nakashima-Sasaki, N. Araki, K. Yatsu, M. Fujita, and S. Umemura. 2016. An isoform of Nedd4-2 is critically involved in the renal adaptation to high salt intake in mice. Sci. Rep. 6:27137. doi:10.1038/srep27137.

49. Molotkov, D., S. Zobova, J.M. Arcas, and L. Khiroug. 2013. Calcium-induced outgrowth of astrocytic peripheral processes requires actin binding by Profilin-1. Cell Calcium. 53:338–348. doi:10.1016/j.ceca.2013.03.001.

50. Mukhopadhyay, D., and H. Riezman. 2007. Proteasome-independent functions of ubiquitin in endocytosis and signaling. Science. 315:201–205. doi:10.1126/science.1127085.

51. Na, C.H., D.R. Jones, Y. Yang, X. Wang, Y. Xu, and J. Peng. 2012. Synaptic Protein Ubiquitination in Rat Brain Revealed by Antibody-based Ubiquitome Analysis. J. Proteome Res. 11:4722–4732. doi:10.1021/pr300536k.

52. Nagaki, K., H. Yamamura, S. Shimada, T. Saito, S. Hisanaga, M. Taoka, T. Isobe, and T. Ichimura. 2006. 14-3-3 Mediates phosphorylation-dependent inhibition of the interaction between the ubiquitin E3 ligase Nedd4-2 and epithelial Na+ channels. Biochemistry. 45:6733–6740. doi:10.1021/bi052640q.

53. Nguyen, Q.-A., M.E. Horn, and R.A. Nicoll. 2016. Distinct roles for extracellular and intracellular domains in neuroligin function at inhibitory synapses. eLife. 5. doi:10.7554/eLife.19236.

54. Oberheim, N.A., X. Wang, S. Goldman, and M. Nedergaard. 2006. Astrocytic complexity distinguishes the human brain. Trends Neurosci. 29:547–553. doi:10.1016/j.tins.2006.08.004.

55. Parente, D.J., C. Garriga, B. Baskin, G. Douglas, M.T. Cho, G.C. Araujo, and M. Shinawi. 2017. Neuroligin 2 nonsense variant associated with anxiety, autism, intellectual disability, hyperphagia, and obesity. Am. J. Med. Genet. A. 173:213–216. doi:10.1002/ajmg.a.37977.

56. Perez-Riverol, Y., J. Bai, C. Bandla, D. García-Seisdedos, S. Hewapathirana, S. Kamatchinathan, D.J. Kundu, A. Prakash, A. Frericks-Zipper, M. Eisenacher, M. Walzer, S. Wang, A. Brazma, and J.A. Vizcaíno. 2022. The PRIDE database resources in 2022: a hub for mass spectrometry-based proteomics evidences. Nucleic Acids Res. 50:D543–D552. doi:10.1093/nar/gkab1038.

57. Pirozzi, G., S.J. McConnell, A.J. Uveges, J.M. Carter, A.B. Sparks, B.K. Kay, and D.M. Fowlkes. 1997. Identification of novel human WW domain-containing proteins by cloning of ligand targets. J. Biol. Chem. 272:14611–14616. doi:10.1074/jbc.272.23.14611.

58. Poulopoulos, A., G. Aramuni, G. Meyer, T. Soykan, M. Hoon, T. Papadopoulos, M. Zhang, I. Paarmann, C. Fuchs, K. Harvey, P. Jedlicka, S.W. Schwarzacher, H. Betz, R.J. Harvey, N. Brose, W. Zhang, and F. Varoqueaux. 2009. Neuroligin 2 Drives Postsynaptic Assembly at Perisomatic Inhibitory Synapses through Gephyrin and Collybistin. Neuron. 63:628–642. doi:10.1016/j.neuron.2009.08.023.

59. Poulopoulos, A., T. Soykan, L.P. Tuffy, M. Hammer, F. Varoqueaux, and N. Brose. 2012. Homodimerization and isoform-specific heterodimerization of neuroligins. Biochem. J. 446:321–330. doi:10.1042/BJ20120808.

60. Preman, P., M. Alfonso-Triguero, E. Alberdi, A. Verkhratsky, and A.M. Arranz. 2021. Astrocytes in Alzheimer’s Disease: Pathological Significance and Molecular Pathways. Cells. 10:540. doi:10.3390/cells10030540.

61. Rodrigues, D.C., M. Mufteev, R.J. Weatheritt, U. Djuric, K.C.H. Ha, P.J. Ross, W. Wei, A. Piekna, M.A. Sartori, L. Byres, R.S.F. Mok, K. Zaslavsky, P. Pasceri, P. Diamandis, Q. Morris, B.J. Blencowe, and J. Ellis. 2020. Shifts in Ribosome Engagement Impact Key Gene Sets in Neurodevelopment and Ubiquitination in Rett Syndrome. Cell Rep. 30:4179–4196.e11. doi:10.1016/j.celrep.2020.02.107.

62. Rothstein, J.D., M. Van Kammen, A.I. Levey, L.J. Martin, and R.W. Kuncl. 1995. Selective loss of glial glutamate transporter GLT-1 in amyotrophic lateral sclerosis. Ann. Neurol. 38:73–84. doi:10.1002/ana.410380114.

63. Roux, K.J., D.I. Kim, M. Raida, and B. Burke. 2012. A promiscuous biotin ligase fusion protein identifies proximal and interacting proteins in mammalian cells. J. Cell Biol. 196:801–810. doi:10.1083/jcb.201112098.

64. Savage, J.T., J.J. Ramirez, W.C. Risher, Y. Wang, D. Irala, and C. Eroglu. 2024. SynBot is an open-source image analysis software for automated quantification of synapses. *Cell Rep*. Methods. 4:100861. doi:10.1016/j.crmeth.2024.100861.

65. Scheiffele, P., J. Fan, J. Choih, R. Fetter, and T. Serafini. 2000. Neuroligin expressed in nonneuronal cells triggers presynaptic development in contacting axons. Cell. 101:657–669.

66. Schnell, E., A.L. Bensen, E.K. Washburn, and G.L. Westbrook. 2012. Neuroligin-1 overexpression in newborn granule cells in vivo. PloS One. 7:e48045. doi:10.1371/journal.pone.0048045.

67. Schnell, E., T.H. Long, A.L. Bensen, E.K. Washburn, and G.L. Westbrook. 2014. Neuroligin-1 knockdown reduces survival of adult-generated newborn hippocampal neurons. Front. Neurosci. 8.

68. Shipman, S.L., and R.A. Nicoll. 2012. Dimerization of postsynaptic neuroligin drives synaptic assembly via transsynaptic clustering of neurexin. Proc. Natl. Acad. Sci. 109:19432–19437. doi:10.1073/pnas.1217633109.

69. Shipman, S.L., E. Schnell, T. Hirai, B.-S. Chen, K.W. Roche, and R.A. Nicoll. 2011. Functional dependence of neuroligin on a new non-PDZ intracellular domain. Nat. Neurosci. 14:718–726. doi:10.1038/nn.2825.

70. Singh, S.K., J.A. Stogsdill, N.S. Pulimood, H. Dingsdale, Y.H. Kim, L.-J. Pilaz, I.H. Kim, A.C. Manhaes, W.S. Rodrigues Jr., A. Pamukcu, E. Enustun, Z. Ertuz, P. Scheiffele, S.H. Soderling, D.L. Silver, R.-R. Ji, A.E. Medina, and C. Eroglu. 2016. Astrocytes Assemble Thalamocortical Synapses by Bridging NRX1α and NL1 via Hevin. Cell. 164:183–196. doi:10.1016/j.cell.2015.11.034.

71. Soto, J.S., Y. Jami-Alahmadi, J. Chacon, S.L. Moye, B. Diaz-Castro, J.A. Wohlschlegel, and B.S. Khakh. 2023. Astrocyte-neuron subproteomes and obsessive-compulsive disorder mechanisms. Nature. 616:764–773. doi:10.1038/s41586-023-05927-7.

72. Stogsdill, J.A., J. Ramirez, D. Liu, Y.H. Kim, K.T. Baldwin, E. Enustun, T. Ejikeme, R.-R. Ji, and C. Eroglu. 2017. Astrocytic neuroligins control astrocyte morphogenesis and synaptogenesis. Nature. 551:192–197. doi:10.1038/nature24638.

73. Sudol, M., and T. Hunter. 2000. NeW Wrinkles for an Old Domain. Cell. 103:1001–1004. doi:10.1016/S0092-8674(00)00203-8.

74. Sudol, M., K. Sliwa, and T. Russo. 2001. Functions of WW domains in the nucleus. FEBS Lett. 490:190–195.

75. Sumita, K., Y. Sato, J. Iida, A. Kawata, M. Hamano, S. Hirabayashi, K. Ohno, E. Peles, and Y. Hata. 2007. Synaptic scaffolding molecule (S-SCAM) membrane-associated guanylate kinase with inverted organization (MAGI)-2 is associated with cell adhesion molecules at inhibitory synapses in rat hippocampal neurons. J. Neurochem. 100:154–166. doi:10.1111/j.1471-4159.2006.04170.x.

76. Tan, C.X., D.S. Bindu, E.J. Hardin, K. Sakers, R. Baumert, J.J. Ramirez, J.T. Savage, and C. Eroglu. 2023. δ-Catenin controls astrocyte morphogenesis via layer-specific astrocyte-neuron cadherin interactions. J. Cell Biol. 222:e202303138. doi:10.1083/jcb.202303138.

77. Tian, R., X. Wu, T.L. Hagemann, A.A. Sosunov, A. Messing, G.M. McKhann, and J.E. Goldman. 2010. Alexander Disease Mutant Glial Fibrillary Acidic Protein Compromises Glutamate Transport in Astrocytes. J. Neuropathol. Exp. Neurol. 69:335–345. doi:10.1097/NEN.0b013e3181d3cb52.

78. Totland, M.Z., C.H. Bergsland, T.A. Fykerud, L.M. Knudsen, N.L. Rasmussen, P.W. Eide, Z. Yohannes, V. Sørensen, A. Brech, R.A. Lothe, and E. Leithe. 2017. E3 ubiquitin ligase NEDD4 induces endocytosis and lysosomal sorting of connexin43 to promote loss of gap junctions. J. Cell Sci. jcs.202408. doi:10.1242/jcs.202408.

79. Varoqueaux, F., G. Aramuni, R.L. Rawson, R. Mohrmann, M. Missler, K. Gottmann, W. Zhang, T.C. Südhof, and N. Brose. 2006. Neuroligins determine synapse maturation and function. Neuron. 51:741–754. doi:10.1016/j.neuron.2006.09.003.

80. Venkatesh, H.S., T.B. Johung, V. Caretti, A. Noll, Y. Tang, S. Nagaraja, E.M. Gibson, C.W. Mount, J. Polepalli, S.S. Mitra, P.J. Woo, R.C. Malenka, H. Vogel, M. Bredel, P. Mallick, and M. Monje. 2015. Neuronal Activity Promotes Glioma Growth through Neuroligin-3 Secretion. Cell. 161:803–816. doi:10.1016/j.cell.2015.04.012.

81. Venkatesh, H.S., L.T. Tam, P.J. Woo, J. Lennon, S. Nagaraja, S.M. Gillespie, J. Ni, D.Y. Duveau, P.J. Morris, J.J. Zhao, C.J. Thomas, and M. Monje. 2017. Targeting neuronal activity-regulated neuroligin-3 dependency in high-grade glioma. Nature. 549:533–537. doi:10.1038/nature24014.

82. Verkhratsky, A., and M. Nedergaard. 2018. Physiology of Astroglia. Physiol. Rev. 98:239–389. doi:10.1152/physrev.00042.2016.

83. Wagner, S.A., P. Beli, B.T. Weinert, C. Schölz, C.D. Kelstrup, C. Young, M.L. Nielsen, J.V. Olsen, C. Brakebusch, and C. Choudhary. 2012. Proteomic analyses reveal divergent ubiquitylation site patterns in murine tissues. Mol. Cell. Proteomics MCP. 11:1578–1585. doi:10.1074/mcp.M112.017905.

84. Wang, S., R. Baumert, G. Séjourné, D.S. Bindu, K. Dimond, K. Sakers, L. Vazquez, J.L. Moore, C.X. Tan, T. Takano, M.P. Rodriguez, N. Brose, L. Bradley, R. Lessing, S.H. Soderling, A.R. La Spada, and C. Eroglu. 2025. PD-linked LRRK2 G2019S mutation impairs astrocyte morphology and synapse maintenance via ERM hyperphosphorylation. BioRxiv Prepr. Serv. Biol. 2023.04.09.536178. doi:10.1101/2023.04.09.536178.

85. Wilson, M.D., S. Sethi, P.J. Lein, and K.P. Keil. 2017. Valid statistical approaches for analyzing sholl data: Mixed effects versus simple linear models. J. Neurosci. Methods. 279:33–43. doi:10.1016/j.jneumeth.2017.01.003.

86. Wu, T., E. Hu, S. Xu, M. Chen, P. Guo, Z. Dai, T. Feng, L. Zhou, W. Tang, L. Zhan, X. Fu, S. Liu, X. Bo, and G. Yu. 2021. clusterProfiler 4.0: A universal enrichment tool for interpreting omics data. The Innovation. 2:100141. doi:10.1016/j.xinn.2021.100141.

87. Xie, H., S. Liu, Y. Fu, Q. Cheng, P. Wang, C.-L. Bi, R. Wang, M.-M. Chen, and M. Fang. 2023. Nuclear access of DNlg3 c-terminal fragment and its function in regulating innate immune response genes. Biochem. Biophys. Res. Commun. 641:93–101. doi:10.1016/j.bbrc.2022.12.030.

88. Xie, Y., A.T. Kuan, W. Wang, Z.T. Herbert, O. Mosto, O. Olukoya, M. Adam, S. Vu, M. Kim, D. Tran, N. Gómez, C. Charpentier, I. Sorour, T.E. Lacey, M.Y. Tolstorukov, B.L. Sabatini, W.-C.A. Lee, and C.C. Harwell. 2022. Astrocyte-neuron crosstalk through Hedgehog signaling mediates cortical synapse development. Cell Rep. 38:110416. doi:10.1016/j.celrep.2022.110416.

89. Xu, C., C.D. Fan, and X. Wang. 2015. Regulation of Mdm2 protein stability and the p53 response by NEDD4-1 E3 ligase. Oncogene. 34:281–289. doi:10.1038/onc.2013.557.

90. Xu, J., Y. Du, J. Xu, X. Hu, L. Gu, X. Li, P. Hu, T. Liao, Q. Xia, Q. Sun, L. Shi, J. Luo, J. Xia, Z. Wang, and J. Xu. 2019. Neuroligin 3 Regulates Dendritic Outgrowth by Modulating Akt/mTOR Signaling. Front. Cell. Neurosci. 13:518. doi:10.3389/fncel.2019.00518.

91. Xu, N., R. Cao, S.-Y. Chen, X.-Z. Gou, B. Wang, H.-M. Luo, F. Gao, and A.-H. Tang. 2024. Structural and functional reorganization of inhibitory synapses by activity-dependent cleavage of neuroligin-2. Proc. Natl. Acad. Sci. 121:e2314541121. doi:10.1073/pnas.2314541121.

92. Zhang, Y., K. Chen, S.A. Sloan, M.L. Bennett, A.R. Scholze, S. O’Keeffe, H.P. Phatnani, P. Guarnieri, C. Caneda, N. Ruderisch, S. Deng, S.A. Liddelow, C. Zhang, R. Daneman, T. Maniatis, B.A. Barres, and J.Q. Wu. 2014. An RNA-Sequencing Transcriptome and Splicing Database of Glia, Neurons, and Vascular Cells of the Cerebral Cortex. J. Neurosci. 34:11929–11947. doi:10.1523/JNEUROSCI.1860-14.2014.

93. Zhang, Y., X. He, X. Meng, X. Wu, H. Tong, X. Zhang, and S. Qu. 2017. Regulation of glutamate transporter trafficking by Nedd4-2 in a Parkinson’s disease model. Cell Death Dis. 8:e2574. doi:10.1038/cddis.2016.454.

94. Zhou, R., S.V. Patel, and P.M. Snyder. 2007. Nedd4-2 catalyzes ubiquitination and degradation of cell surface ENaC. J. Biol. Chem. 282:20207–20212. doi:10.1074/jbc.M611329200.

