## Supplementary material for "Neuroligin-2 is ubiquitinated by Nedd4l to control developmental astrocyte morphogenesis": Table S1

| **Figure Panel** | **Description** | **Statistical Analysis** |
| --- | --- | --- |
| 1D | Sholl analysis | Linear mixed effects model (LME) ANOVA F(3, 319) = 42.46, p < 2.2E-16. Dunnett’s posthoc tests values on graph [control condition is shNL2+HA-NL2-RM]. N = 48-96 cells across conditions, 3 biological replicates. Data distribution was assumed to be normal but this was not formally tested. |
| 1F | Sholl analysis | Linear mixed effects model (LME) ANOVA F(3, 311)=15.5, p=2.02E-9. Dunnett’s posthoc tests values on graph [control condition is shCtrl+HA-NL2-RM]. N = 53-97 cells across conditions, 3 biological replicates. Data distribution was assumed to be normal but this was not formally tested. |
| 2C | Sholl analysis | Linear mixed effects model (LME) ANOVA F(3, 159)=1.97, p=0.12. Dunnett’s posthoc tests values on graph [control condition is shNL2+HA-NL2-RM]. N = 36-46 cells across conditions, 3 biological replicates. Data distribution was assumed to be normal but this was not formally tested. |
| 2F | Sholl analysis | Linear mixed effects model (LME) ANOVA F(3,356)=38.22, p<2.2E-16. Dunnett’s posthoc tests values on graph [control condition is shNL2+HA-NL2-RM]. N = 57-97 cells across conditions, 4 biological replicates. Data distribution was assumed to be normal but this was not formally tested. |
| 2J | Sholl analysis | Linear mixed effects model (LME) ANOVA F(3, 304)=25.76, p=7.1E-15. Dunnett’s posthoc tests values on graph [control condition is shNL2+HA-NL2-RM]. N = 71-84 cells across conditions, 4 biological replicates. Data distribution was assumed to be normal but this was not formally tested. |
| 4A | Overlap of BioID candidates with previously published datasets | Fisher’s exact test. Exact p-values: Kang et al [Astrocyte NL2 BioID] = 1.4e-40 Kang et al [Neuron NL2 BioID] = 3.5e-30  Loh et al [Astrocyte NL2 BioID] = 4.4e-6 Loh et al [Neuron NL2 BioID] = 7.1e-18  Poulopoulos et al [Astrocyte NL2 BioID] = 1.4e-13 Poulopoulos et al [Neuron NL2 BioID] = 3.9e-13 |
| 4B | BioID enrichment | Heteroscedastic t-test. P-values printed on graph are uncorrected p-values. |
| 4C | Gene ontology [Astrocyte NL BioID comparison] | Over-representation analysis with FDR correction. Used clusterProfiler package (v4.16.0) compareCluster function. |
| 4D | Gene ontology | Over-representation analysis with FDR correction. Used clusterProfiler package (v4.16.0) compareCluster function. |
| 4E | BioID enrichment | Heteroscedastic t-test. P-values printed on graph are uncorrected p-values. Fold change is relative to cell-type specific soluble BirA control. Values come from Table S1. |
| 5B | Sholl Analysis | Linear mixed effects model (LME) ANOVA F(3, 332)=217.36, p<2.2E-16. Tukey’s posthoc tests values on graph [control condition is shCtrl]. N = 60-100 cells across conditions, 4 biological replicates. Data distribution was assumed to be normal but this was not formally tested. |
| 5E | Territory volume analysis | Nested ANOVA F(1,8) = 11.95, p=0.0023. 4-7 animals imaged per condition, with at least 2 images per animal acquired. |
| 5F | Sholl analysis | Linear mixed effects model (LME) ANOVA F(1, 9) = 5.59, p=0.04. 4-7 animals imaged per condition, with at least 2 images per animal acquired. Data distribution was assumed to be normal but this was not formally tested. |
| 6C | Densitometry quantification of I.P. bands | Paired student’s t-test. P=0.103. N=4 independent replicates per condition. Equal variance was tested using var.test() in R. |
| 6E | Densitometry quantification of I.P. bands | Paired student’s t-test. P=0.034. N=3 independent replicates per condition. Equal variance was tested using var.test() in R. |
| 6G | Densitometry quantification of I.P. bands | Paired student’s t-test. P=0.043. N=3 independent replicates per condition. Equal variance was tested using var.test() in R. |
| 6I | Densitometry quantification of I.P. bands | Paired student’s t-test. P=0.006. N=3 independent replicates per condition. Equal variance was tested using var.test() in R. |
| 7B | Quantification of high molecular weight HA-NL2 [> 100kDa] | Paired student’s t-test. P=0.04. N=3 independent replicates per condition. Data distribution was assumed to be normal but this was not formally tested. |
| 7D | Densitometry quantification of HA-NL2 western blots | Two-way ANOVA: Condition: F(1,16) = 15.018, p=0.00134 Timepoint: F(3,16) = 0.41, p=0.748 Interaction Condition:Timepoint: F(3, 16) = 2.242, p=0.123. Data distribution was assumed to be normal but this was not formally tested. |
| 7F | Densitometry quantification of HA-NL2 western blots | Two-way ANOVA: Condition: F(1,16) = 1.392, p=0.255 Timepoint: F(3,16) = 1.681, p=0.211 Interaction Condition:Timepoint: F(3, 16) = 2.328, p=0.113. Data distribution was assumed to be normal but this was not formally tested. |
| 8B | Sholl analysis | Linear mixed effects model (LME) ANOVA F(3, 284)=103.14, p<2.2E-16. Dunnett’s posthoc tests values on graph [control condition is shCtrl]. N = 62-80 cells across conditions, 3 biological replicates. Data distribution was assumed to be normal but this was not formally tested. |
| 8D | Territory volume | Kruskal-Wallis rank sum test. χ2 = 19.5, df=3, p=0.00013. Wilcoxon rank sum post-hoc tests with Holm-Šídák multiple comparison correction are on the graph. Data was tested for normality using a quantile-quantile plot in R. |
| 8E | Sholl analysis | Linear mixed effects model (LME) ANOVA F(3, 102)=5.08, p=0.0025. Tukey’s posthoc results in figure. N = 21-33 cells total from 6-7 animals per group, with at least 2 cells imaged per animals. Data distribution was assumed to be normal but this was not formally tested. |
| S1A | Densitometry quantification of I.P. bands | Student’s t-test. P=0.017 [WT NL2 shCtrl vs shNL2]; P=0.61 [RM NL2 shCtrl vs shNL2]. N=3 independent replicates per condition. Equal variance was tested using var.test() in R. |
| S1B | Densitometry quantification of I.P. bands | Student’s t-test. P=0.124 [NL1: shCtrl vs shNL2]; P=0.026 [NL2: shCtrl vs shNL2]; P=0.125 [NL3: shCtrl vs shNL2]. N=3 independent replicates per condition. Equal variance was tested using var.test() in R. |
| S1D | Sholl analysis | Linear mixed effects model (LME) ANOVA F(3, 286)=75.01, p<2.2E-16. Dunnett’s posthoc tests values on graph [control condition is shCtrl]. N = 53-96 cells across conditions, 3 biological replicates. Data distribution was assumed to be normal but this was not formally tested. |
| S1H | Sholl analysis | Linear mixed effects model (LME) ANOVA F(2, 6) = 21.12, p=0.0019. Tukey’s posthoc tests values on graph [control condition is shCtrl]. N = 40 cells across conditions, 3 biological replicates. Data distribution was assumed to be normal but this was not formally tested. |
| S3C | Sholl analysis | Linear mixed effects model (LME) ANOVA F(1, 3) = 11.01, p=1.4E-6. Dunnett’s posthoc tests values on graph [control condition is shNL2+HA-NL2-RM]. N = 14-56 cells across conditions, 3 biological replicates. Data distribution was assumed to be normal but this was not formally tested. |
| S4A-F | BioID enrichment. | Heteroscedastic t-test. P-values printed on graph are uncorrected p-values. Fold change is relative to cell-type specific soluble BirA control. Values come from Table S1. |
| S5A | Densitometry of Nedd4l western blot, normalized to Actin | Student’s t-test. P-value = 0.0004. N=3 independent replicates. Data distribution was assumed to be normal but this was not formally tested. |
| S5F | Quantification of excitatory and inhibitory synapse density in shNedd4l or shCtrl astrocytes | Linear mixed effects ANOVA. Excitatory synapses: F(1,13) = 0.12, p=0.73 Inhibitory synapses: F(1,13) = 0.61, p=0.45  N = 8 shCtrl animals, and 7 shNedd4l animals. 4-8 images per animal were acquired. Data distribution was assumed to be normal but this was not formally tested. |
